# Mitochondrial and protein homeostasis pathways are transcriptionally impaired in islets during type 1 diabetes pathogenesis

**DOI:** 10.64898/2026.07.29.741469

**Authors:** Alexandra E. Cuaycal, Elizabeth A. Butterworth, Scott Stimpson, Jing Chen, Nataliya I. Lenchik, Leigh A. Baratta, Edward A. Phelps, Scott Grieshaber, Mark A. Atkinson, Wei-Jun Qian, Martha Campbell-Thompson, Ivan C. Gerling, Clayton E. Mathews

**Affiliations:** Department of Infectious Diseases and Immunology, University of Florida, Gainesville, FL, USA; J. Crayton Pruitt Family Department of Biomedical Engineering, University of Florida, Gainesville, FL, USA; Department of Pathology, Immunology and Laboratory Medicine, University of Florida, Gainesville, FL, USA; School of Medical Sciences, Faculty of Medicine and Health, The University of Sydney, Sydney, NSW, Australia; Department of Medicine, University of Tennessee, Memphis, TN, USA; Department of Biological Sciences, University of Idaho, Moscow, ID, USA; Biological Sciences Division, Pacific Northwest National Laboratory, Richland, WA, USA

## Abstract

The decline in first-phase insulin response (FPIR) during the presymptomatic period of type 1 diabetes (T1D) is well established. *In-situ* functional studies with pancreas tissue slices showed that β-cell loss of glucose-responsiveness was independent of T-cell infiltration into islets in recent-onset T1D cases. However, the mechanisms driving β-cell dysfunction before the onset of T1D remain unclear. In pancreas tissue from donors across the natural history of T1D, we utilized an *in-situ,* whole*-*islet phenotypical and transcriptomic approach to unravel novel targets in the glucose-stimulus coupled secretion pathway that are similarly impaired in T-cell infiltrated and non-infiltrated islets. Specifically, we observed that islets from autoantibody positive (single(s) or multiple(m) AAb+) donors exhibited activation of post-transcriptional gene regulation along with reduced protein translation, processing in the endoplasmic reticulum (ER), and ER stress. Disrupted mitochondrial metabolism and bioenergetics were prominent in islets from multiple AAb+ and T1D donors with disease durations ≤7 years. In addition, T1D islets presented reduced mitochondrial protein import, quality control, and dynamics, together with downregulated genes in insulin secretory pathways. During infiltration, these pathways remain dysregulated while immune/inflammatory transcripts were increased. These studies identified novel mechanisms of β-cell dysregulation before symptomatic onset and independent of T-cell infiltration in T1D pathogenesis.

## Introduction

Type 1 Diabetes (T1D) is an autoimmune disease considered to result from direct T-cell-mediated destruction of insulin-producing pancreatic β-cells (1,2). The pathogenesis and natural history of T1D is not fully understood but typically includes the development of autoantibodies (AAbs) against novel β-cell antigens followed by a progressive decline in first-phase insulin response (FPIR) (3–7). Recent studies indicate that reduced FPIR can be evident as early as 4-6 years prior to disease onset and drastically declines within the last two years before T1D diagnosis (5,8,9). This β-cell impairment results in disrupted glucose homeostasis, which, together with β-cell loss, defines the onset of T1D (2,10). The loss of FPIR has not only been observed in clinical studies but also *in situ* with the pancreas tissue slice platform (11,12). Importantly, our previous work with live imaging of T1D pancreatic slices revealed that this loss of glucose-responsiveness occurs irrespective of the presence of CD3+ T-cells (11). These data call into question the direct impact of T-cells on β-cell dysfunction and highlight an urgent need to define the mechanisms driving β-cell failure and disrupted glucose homeostasis prior to the onset of T1D.

As *in-situ* longitudinal studies of human islet cell biology are impossible in living subjects due to ethical reasons, we used pancreas tissues obtained from the Network for Pancreatic Organ donors with Diabetes (nPOD) program to phenotypically and transcriptionally characterize multiple islets across organ donors during the natural history of T1D (13) (Supplemental Figure 1). We analyzed serial, frozen pancreas sections encompassing entire islets for both immunostaining (immune, mitochondrial, and β-cell markers) and laser microdissection (LMD) with transcriptomic analysis. LMD of preserved islets allowed for analysis of changes across study subject groups (No diabetes: ND, single (s)AAb+, multiple (m)AAb+ and T1D) that could be altered during standard islet isolation and dissociation processes (14,15). Using this approach, we investigated transcriptomic patterns of islet dysfunction and stress, based on β-cell and CD3+ cell infiltration status.

Specifically, we characterized transcriptional changes in islets across the natural history of T1D to provide mechanistic evidence underlying the loss of FPIR and to define changes at the islet level that were independent and dependent on local autoimmunity (i.e., insulitis). We hypothesized that β-cells lose FPIR through dysregulated processes in the glucose stimulus secretion coupling pathway, independent of current, local T-cell infiltration. Herein, we provide a unique and extensive *in-situ* analysis of human islet cell biology over the natural history of T1D including the transcriptional signatures of islets with and without CD3+ T-cell infiltration. By comparing non-infiltrated islets from donors with no diabetes versus those from donors without T1D that have seroconverted (i.e., AAb+), or donors with T1D and insulin positive (INS+) islets (< 10 years duration), novel gene signatures were identified in islet cell bioenergetics, mitochondrial, and protein homeostasis, as well as insulin secretory pathways that are impaired in T1D pathogenesis, irrespective of local insulitis.

## Results

### Single islets were successfully microdissected from four groups of donors

Donors were selected based on INS+ islets and insulitis status during initial histopathology reviews (10). A single non-diabetic (ND) islet studied had ≥6 CD3+ cells/islet. Based on previously defined consensus criteria (16), this donor did not have insulitis. One of six (17%) sAAb+ and two of five (40%) mAAb+ donors had insulitis (**Supplemental Table 1).** All 17 donors with T1D had islets with residual β-cells (INS+), with and without insulitis (**Supplemental Table 1**).

Donor sex, age, and BMI were similar across groups (**Figure 1A-C**, **Supplemental Table 1**) except the inclusion of one 69-year-old mAAb+ case resulted in significant age differences in mAAb+ vs. ND and T1D, respectively (**Figure 1A-B**). BMI was similar across the groups (Figure 1C). Pancreas weights were, as expected, significantly lower in T1D cases when normalized to body weight (relative pancreas weight, RPW) (**Figure 1D**) (17). Duration of diabetes ranged from 0 to 7 years (2.6 ± 2.2 years), with four of 16 cases (25%) having recent-onset T1D (˂1 year disease duration) (**Supplemental Table 1**).

**Figure 1.**
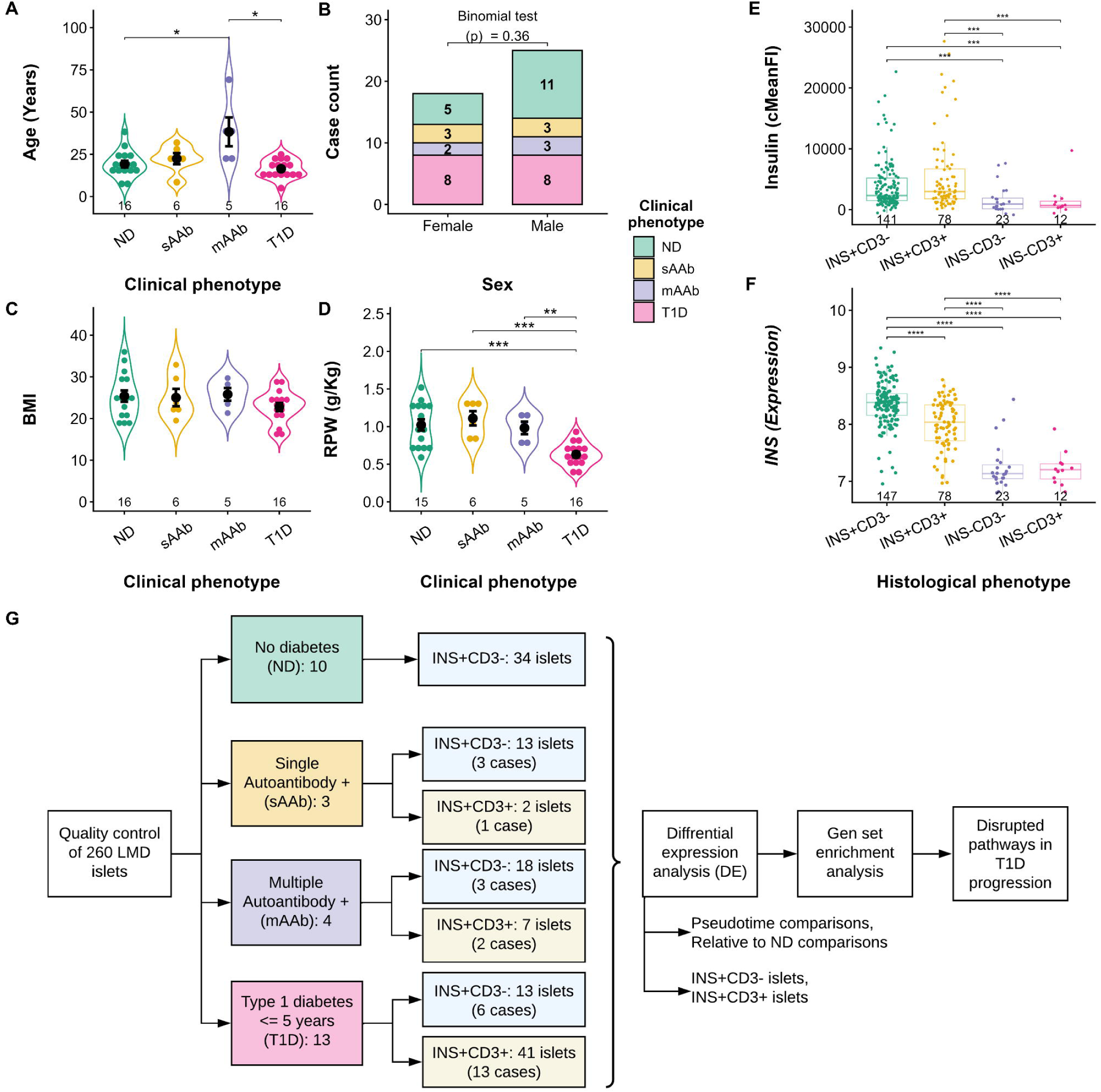
Donor and islet characteristics. Donor cohort age **(A)**, sex **(B)**, body-mass index (BMI) **(C)** and relative pancreas weight (RPW) **(D)** distribution. **(E)** Quantification of islet insulin by immunofluorescence staining of human pancreas sections and **(F)** same-islet *INS* gene expression across the islet histological phenotypes. The immunofluorescence signal was corrected by subtracting the acinar tissue signal from the islet region (cMeanFI). **(G)** Flow chart of bioinformatics analysis of the final islet set after quality control. Statistical significance of donor characteristics and cMeanFI was assessed by using the non-parametric Wilcoxon test, except for donor’s sex (binomial test). Gene expression differences in F were assessed with a one-way ANOVA followed by Tukey post-hoc. Multiple testing correction was performed with the Benjamini-Hochberg (BH) method. Adjusted *p*: ∗*p*<0.05, ∗∗*p*<0.01, ∗∗∗*p*<0.001, ∗∗∗∗*p*<0.0001.

260 LMD islets were obtained across donor groups for subsequent transcriptomics analyses (**Supplemental Figure 1A**), including 147 INS+CD3-, 78 INS+CD3+, 23 INS-CD3-, and 12 INS-CD3+ islets as determined by immunofluorescence (**Figure 1E, Supplemental Figure 1B, Supplemental Table 2**). The INS corrected mean fluorescence intensity (cMeanFI) was similar between INS+CD3- and INS+CD3+ islets, with INS-islets having no visible insulin staining (**Figure 1E**). Similarly, the *INS* gene expression was significantly lower in INS-islets, validating the *a priori* immunohistological islet phenotyping strategy (**Figure 1F**). The mean RNA Integrity Number (RIN) number for the entire 260 islet set was 5.8 ± 1.2, and histological groups within each clinical group also had comparable RIN values with average RIN > 5 (**Supplemental Figure 1C**).

### Quality control and exploratory analysis of 260 islet transcriptomics

Quality control (QC) check with Uniform Manifold Approximation and Projection (UMAP) demonstrated good clustering of islets across the four clinical phenotypes (**Supplemental Figure 2A**). T1D islets clustered together irrespective of their histological phenotype (**Supplemental Figure 2B**). Initial differential expression (DE) analysis with INS+CD3-islets revealed non-β-cell transcripts (expected to be present) in our dataset. Hence, we utilized cell-type-specific markers from scRNA-seq of HPAP islets (18) to visualize enrichment/depletion of different cell types in our 260 islets (**Supplemental Figure 2C**). We focused on α-cell contamination, given the relatively high proportion of this cell type within an islet. We calculated z-scores for the β-cell-specific markers across the 260 islets and filtered out *INS* positive, β-cell-containing islets (i.e., INS+ islets) that had z-scores >1 for three or more α-cell-specific transcripts (43 islets), as this indicated an over-representation of α-cells.

We then filtered islets based on the donor HLA Class II genetic risk for T1D. HLA class II was stratified into protective (DR15, DR15/X), neutral, or risk for T1D (DR3, DR4, DR3/4) (19–21). Islets from donors with protective HLA Class II (DR15) were excluded (65 islets). Finally, only INS+ islets (based on histology and gene expression) were used for subsequent analyses (129/152 islets). The final dataset for the analyses described below consisted of 35 ND islets (34 INS+CD3-, 1 INS+CD3+), 15 sAAb+ islets (13 INS+CD3-, 2 INS+CD3+), 25 mAAb+ islets (18 INS+CD3-, 7 INS+CD3+), and 54 T1D islets (13 INS+CD3-, 41 INS+CD3+) (**Figure 1G**, **Supplemental Table 3**). Analyses with INS+CD3+ islets included only the mAAb+ and T1D groups due to the dearth of INS+CD3+ islets in the ND (n=1) and sAAb+ (n=2) groups.

This final islet set had consistent mean RIN > 5 with no significant differences observed across the histological phenotypes within each clinical group (**Supplemental Figure 1D**). There were no significant correlations between housekeeping gene expression and RIN (**Supplemental Figure 3A**). The insulin stain area was not significantly different among clinical groups (**Supplemental Figure 3B**), corroborating the *in-silico* filtering approach. Nonetheless, it was reduced in T1D islets after normalization to the average islet cut area, consistent with reduced β-cell mass (**Supplemental Figure 3C**). We first investigated the transcriptional changes that occur in INS+CD3-islets across the four different clinical phenotypes. Subsequently, INS+CD3- and INS+CD3+ islets from mAAb+ or T1D individuals were compared to describe changes that occur in the absence or presence of current immune cell infiltration. Indeed, CD3E/G and CD68 expression were significantly increased in the INS+CD3+ islets of mAAb+ and T1D persons compared to CD3- islets in these two groups (**Supplemental Figure 4**) (22).

### Islet transcriptional changes occur early in T1D natural history

We first performed DE analysis with INS+CD3- and two different comparisons: each clinical group compared to the ND group and pseudo-time comparisons (ND→sAAb+→mAAb+→T1D) following the natural progression of T1D (**Figure 2A and 2B**). DE was defined with an absolute log-fold change (logFC)>0.5 and an adjusted *p*<0.001. **Supplemental Table 4** displays the overall DE results for all the contrasts evaluated. We performed both over-representation (ORA) and gene set enrichment analyses (GSEA) with gene ontology (GO) biological processes (BP), as well as KEGG pathways for each contrast analyzed, as noted in Methods. **Supplementary Tables** are provided for each analysis as described below.

**Figure 2.**
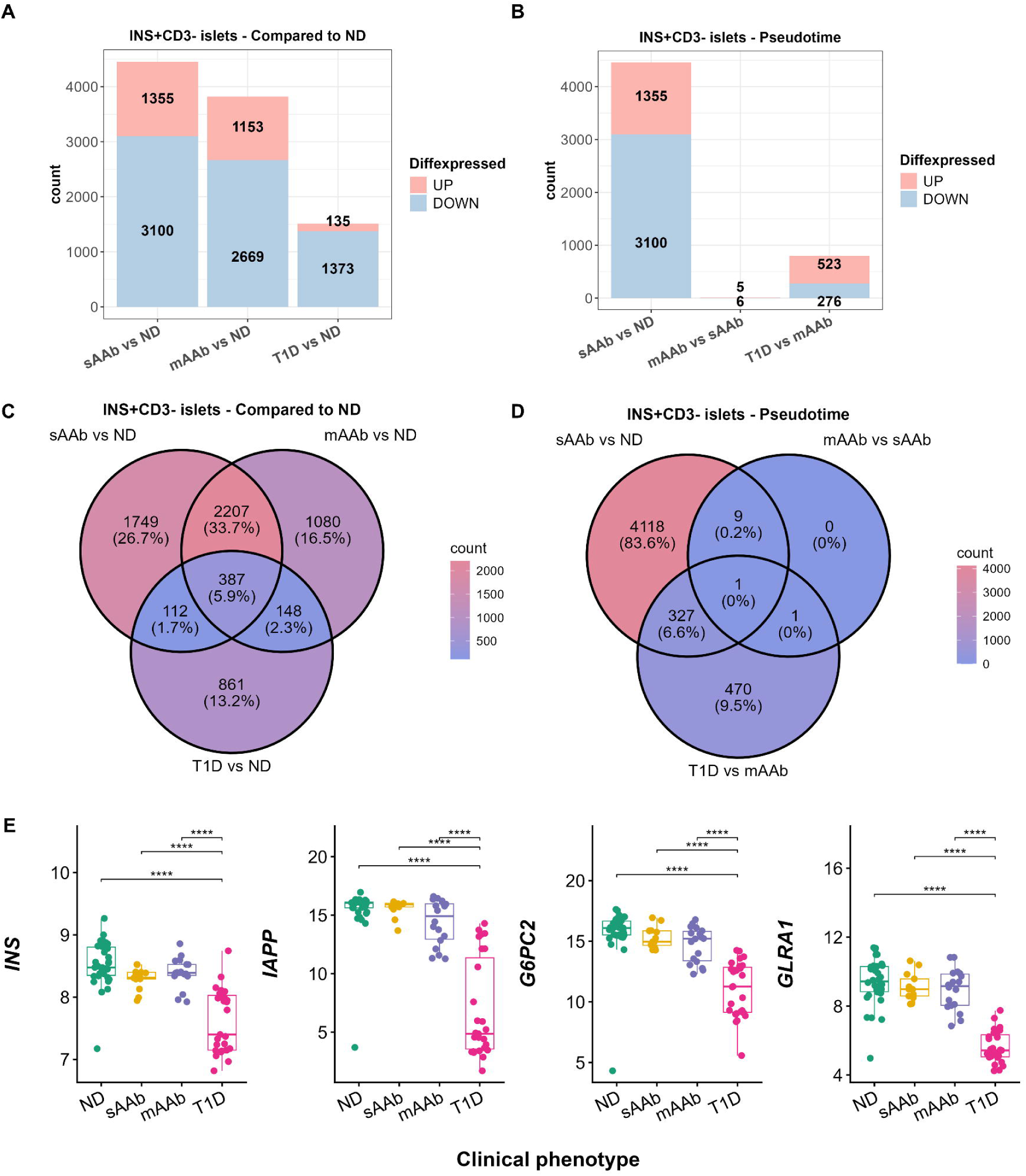
Differentially expressed genes (DEGs) in INS+CD3-islets across the natural history of T1D. **(A)** Number of DEGs in INS+CD3-islets from single(s), multiple(m) AAb positive and T1D cases when compared to no diabetes (ND). **(B)** Number of DEGs in INS+CD3-islets from pseudo-time comparisons (sAAb+ vs. ND, mAAb+ vs. sAAb+ and T1D vs. mAAb). DEGs were defined as genes with an absolute logFC˃0.5 and an adjusted *p*˂0.001. **(C)** Venn diagrams of DEGs across comparisons to ND and **(D)** pseudo-time comparisons. **(E)** β-cell marker gene expression in INS+CD3-islets across clinical phenotypes. Gene expression differences in E were assessed with a one-way ANOVA followed by Tukey post-hoc. Multiple testing correction was performed with the Benjamini-Hochberg (BH) method. Adjusted *p*: ∗*p*<0.05, ∗∗*p*<0.01, ∗∗∗*p*<0.001, ∗∗∗∗*p*<0.0001.

We observed 4,455 DEG in sAAb+ islets, with the majority (3,100 DEG, ∼70%) being downregulated compared to ND persons (**Figure 2A**). GSEA analysis demonstrated modulation of over 500 BP in sAAb+ subjects (**Supplemental Table 5**) with only one BP being increased: activation of post-transcriptional gene expression regulation, specifically, micro (mi)RNA-mediated gene silencing. Metabolic/bioenergetic pathways were among the most significantly suppressed by KEGG GSEA analysis (**Supplemental Figure 5B**, **Supplemental Table 6**). Additionally, BP that were suppressed include: response to bacterium, humoral immune response, wound healing, and epithelial cell proliferation (**Supplemental Figure 5A**).

Similar results were observed comparing mAAb+ to ND islets (**Figure 2A**): a total of 3,822 DEG with ∼70% reduced in those with mAAb+. KEGG pathways GSEA revealed the suppression of metabolic/bioenergetic pathways as well as the suppression of protein processing in the ER, lysosome, and cellular adhesion pathways (**Supplemental Figure 5D**, **Supplemental Table 7**). As observed in those with sAAb+i, islets from those mAAb+ showed activated post-transcriptional gene silencing and suppressed genes involved in wound healing, regulation of proteolysis, cellular response to stress, generation of precursor metabolites, and energy (**Supplemental Figure 5C**, **Supplemental Table 8**). For pseudo-time analysis, the sAAb+ vs. ND and mAAb+ vs. ND revealed striking similarities between sAAb+ and mAAb+ islets (**Figure 2B**), with over 2,500 DEG shared and in the same direction (**Figure 2C**). These results suggest that the transcriptional changes occurring in sAAb+ islets are maintained in mAAb+ islets. There were only 11 DEG in mAAb+ vs. sAAb+ islets (**Figure 2B**, **Supplemental Table 4**): three corresponded to long non-coding (lnc)RNA (*BCYRN1*, *ENST00000425419*, *ENST00000420548*), one transcriptional repressor (*RIPPLY2*), a protein involved in ribosomal QC (*N4BP2*), and a component of the elongator complex involved in tRNA modifications (*ELP5*). The mitochondrial F1F0 ATP Synthase Membrane Subunit E (*ATP5ME*), while increased in sAAb+ vs. ND persons, was significantly reduced in mAAb+ vs. ND and vs. sAAb+ subjects. Indeed, 10 genes transitioned from being upregulated in sAAb+ (vs. ND) to suppressed in mAAb+ vs. ND individuals. In contrast, *ELP5* was reduced in sAAb+ subjects and elevated in those mAAb+. *N4BP2* was downregulated uniquely in the mAAb+ vs. sAAb+ subject comparison. Finally, the additional four genes corresponded to acinar-cell genes present in our dataset. Since our cohort included a mAAb+ case of a 69 year old individual (nPOD case 6080), we performed additional DE analyses with age-matched cases (ND: 26.4 ± 3.1 years, sAAb+: 25.6 ± 5.7 years, mAAb+: 38.7 ± 2.3 years, T1D: 20.3 ± 4.1 years). Similar processes were DE in this islet subset as in the whole dataset (**Supplemental Figure 6**).

In INS+CD3-islets, the T1D vs. ND comparison yielded over 1,500 DEG (Figure 2A). Most were downregulated with suppressed BP in cell-signaling, regulation of hormone levels, and secretion (**Supplemental Figure 5E**, **Supplemental Table 9**). Importantly, the loss of genes involved in oxidative phosphorylation (OXPHOS) and the proteasome from ND to the AAb+ groups was further enriched in T1D islets (**Supplemental Figure 5F**, **Supplemental Table 10**). Several neurodegenerative disease pathways were noted as significant in GSEA due to the enrichment of OXPHOS genes in those BP. T1D islets downregulated β-cell-specific genes including *IAPP* (logFC: -5.9) and *G6PC2* (logFC: -4.3), consistent with an expected loss of β-cells after T1D onset (1). *GLRA1* was also downregulated (logFC: -3.5) (**Figure 2E**). This gene encodes a glycine receptor previously noted to be specific for β-cells (23). These data were corroborated by the insulin staining area results (**Supplemental Figure 3C**). The pseudo-transition from mAAb+ to T1D revealed over 700 DEG (**Figure 2B**). Conversely to the ND comparison, most of the DEG were upregulated. Indeed, GSEA analysis demonstrated activation of seven BP, including proteolysis, regulation of catalytic activity, response to stress, and immune response in islets of cases with T1D compared to those mAAb+ (**Supplemental Figure 5G**, **Supplemental Table 11**).

### Post-transcriptional gene regulation is activated in sAAb+ and mAAb+ islets

Our pseudotime analysis revealed striking similarities between the sAAb+ and mAAb+ groups. From the 2,594 DEG shared, 2,207 DEG were unique between sAAb+ and mAAb+ persons when compared to those ND (**Figure 2C**). Similar to sAAb+ islets, there were 1,153 DEG that were significantly upregulated in mAAb+ vs. ND islets. GSEA analyses revealed that both sAAb+ and mAAb+ islets activated post-transcriptional gene silencing (**Supplemental Figure 5 A,C**). Among both comparisons, there were 61 enriched non-coding genes, including 60 miRNAs and an uncharacterized lncRNA (**Supplemental Figure 7**). We mined two miRNA-target interaction databases for their target mRNA and compared them to the DEG in sAAb+ and mAAb+ vs. ND individuals. DE miRNA targets were investigated using the prediction-based miRDB database (24) and the experimentally validated database miRTarBase (25). Overall, we retrieved miRNA target data for 47 out of the 60 DE miRNA (**Supplemental Table 12**). We observed the significant enrichment, after multiple testing correction, of miRNA targets for 21 of the 47 miRNAs interrogated (adjusted *p*<0.001, odds ratio (OR)>1). As a control, we performed enrichment analysis for miRNAs that were not significantly DE across the three groups (i.e., sAAb+, mAAb+, and T1D) and expectedly observed non-significant enrichments across all miRNAs evaluated (**Supplemental Table 13**).

Remarkably, the target enrichment for DE miRNAs represented ∼20-50% of the total targets predicted/validated from both databases and present in the background genes in our islet gene expression data (n=31,128 genes). From the miRNAs that had a significant enrichment in both databases, we identified 13 miRNAs with an OR>2 in the miRTarBase database (validated targets) (**Table 1**), indicating a ≥ two-fold increased likelihood of being DEG compared to non-target genes. The top three miRNA with OR> 2 in miRTarBase were hsa-miR-129, hsa-miR-551b, and hsa-miR-1226. Subsequently, we performed ORA for the DE targets identified across both databases for each of these 13 miRNAs (**Supplemental Table 14**). We observed enrichment in genes related to import to nucleus, cytoplasmic translation, protein import to nucleus, ER-to-Golgi vesicle-mediated transport, response to ER stress, cell-substrate adhesion, and cell-matrix adhesion (**Table 1**). Substantiating the importance of these DE miRNAs were the similar biological processes suppressed in sAAb+ and mAAb+ islets (**Supplemental Figure 5A & 5C**) upon analysis of the total RNA.

**Table 1.** Over-representation analysis (ORA) with gene ontology (GO) biological processes (BP) of miRNA targets. ORA was run with the DEGs in the INS+CD3-sAAb+ and mAAb+ islets versus ND comparison that were also significant miRNA-targets with odds ratios over 2 in the miRTarBase database. Significant BPs were considered with an adjusted *p*<0.05. The miRNA enrichments were performed with a Fisher’s exact test and corrected for multiple testing with the Benjamini-Hochberg (BH). Significant miRNA-target enrichments were considered with an adjusted *p˂*0.001 and an OR>1. See Table S12 for the full list of miRNA target enrichment results including the number of background genes and enriched DEGs. The miRNAs presented here are highlighted in blue in Table S12. n.i.: No enriched pathways identified.

| miRNA name | Gene symbol | Ensembl ID | Donor type expression | DEG target GO BP over-representation |
| --- | --- | --- | --- | --- |
| hsa-miR-129 | *MIR129-1* | ENSG00000207705 | sAAb+only | n.i. |
| hsa-miR-551b | *MIR551B* | ENSG00000207717 | sAAb+and mAAb | Response to endoplasmic reticulum stress; regulation of translation; organic cation transport |
| hsa-miR-1226 | *MIR1226* | ENSG00000221585 | mAAb+ only | Positive regulation of protein-containing complex disassembly; regulation of protein-containing complex disassembly; cell-substrate adhesion |
| hsa-miR-1260b | *MIR1260B* | ENSG00000266192 | sAAb+and mAAb | Response to endoplasmic reticulum stress; viral process; peptidyl-threonine modification |
| hsa-miR-142 | *MIR142* | ENSG00000284353 | mAAb+ only | Positive regulation of cell adhesion; endosomal transport; protein localization to vacuole |
| hsa-miR-193a | *MIR193A* | ENSG00000207614 | mAAb+ only | Regulation of protein localization to endosome; protein localization to endosome |
| hsa-miR-877 | *MIR877* | ENSG00000216101 | mAAb+ only | Protein localization to nucleus; nucleocytoplasmic transport; nuclear transport |
| hsa-miR-449b | *MIR449B* | ENSG00000207728 | mAAb+ only | Ameboidal-type cell migration; epithelial cell development; podocyte development |
| hsa-miR-208a | *MIR208A* | ENSG00000199157 | sAAb+and mAAb | Nuclear-transcribed mRNA catabolic process, deadenylation-dependent decay; regulation of cell cycle phase transition; organ growth |
| hsa-miR-149 | *MIR149* | ENSG00000207611 | mAAb+ only | Viral process; regulation of translation; negative regulation of catabolic process |
| hsa-miR-98 | *MIR98* | ENSG00000271886 | mAAb+ only | Cell-substrate adhesion; response to transforming growth factor beta; cell-matrix adhesion |
| hsa-miR-365a | *MIR365A* | ENSG00000199130 | mAAb+ only | Intracellular receptor signaling pathway |
| hsa-miR-33b | *MIR33B* | ENSG00000207839 | sAAb+and mAAb | n.i. |

We also observed the upregulation of four (U1, U2, U4, U6) of the five small nuclear RNAs (snRNAs) that form the spliceosome complex (**Supplemental Table 4**) (26). Notably, the U6 snRNA genes, forming the spliceosome active site, were the most upregulated, with logFCs from 0.5-3 in mAAb+ vs. ND islets (**Supplemental Table 4**). In accordance with this result, the splicing regulator *NOVA1* was also upregulated in sAAb+ islets (logFC=1.3) with a similar trend in mAAb+ islets (logFC=0.5, adjusted *p*=0.07). *NOVA1* has been shown to regulate multiple processes, including insulin secretion (27).

### INS+CD3-sAAb+ islets exhibit downregulation of defense response, extracellular matrix remodeling, and cellular adhesion genes

To identify processes that occur distinctively in sAAb+, mAAb+, and T1D islets, we filtered DEG unique to the sAAb+ vs. ND comparison (1,749 DEG; **Figure 2C**). 239 BPs were enriched by GSEA (adjusted *p*<0.01, **Supplemental Table 15**). The top 15 significantly suppressed BP corresponded mostly associated with immune response and cell adhesion mechanisms (**Figure 3A**). Similar pathways were also observed with the full DEG set for sAAb+ vs. ND comparison (**Supplemental Figure 5A-B**). Nine KEGG pathways were enriched by ORA with adjusted *p*<0.05 (**Supplemental Table 16**). Extracellular matrix (ECM)-receptor interaction, focal adhesion, and ribosome biogenesis in eukaryotes were the top three enriched pathways (**Figure 3B**).

**Figure 3.**
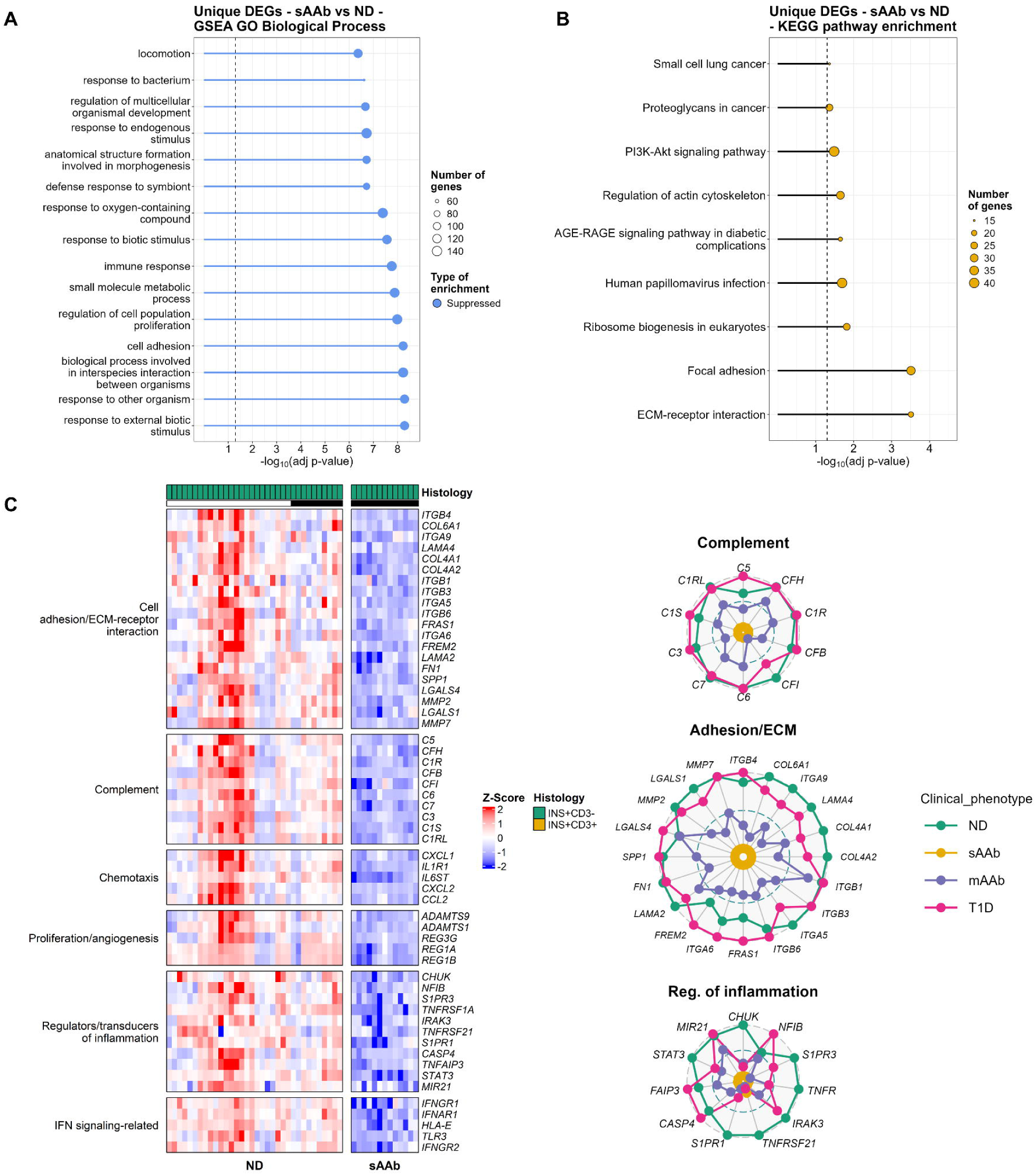
Dysregulated pathways in INS+CD3-islets from sAAb+ positive cases when compared to ND. **(A)** Gene set enrichment analysis (GSEA) of gene ontology (GO) with biological processes (BP) and **(B)** KEGG pathway over-representation analysis (ORA) of unique DEGs in INS+CD3-sAAb+ islets. **(C)** Heatmap of representative DEGs in INS+CD3-sAAb+ islets. Gene expression in the heatmap is presented as z-scores. Black line indicates islets from age-matched subcohort. **(D)** Radial plots of islet average gene expression across clinical groups for representative DEGs in INS+CD3-sAAb+ islets. Gene expression in radial plots is scaled from 0-1, where 0 and 1 represent the minimum and maximum average gene expression values, respectively. Statistical significance for BPs and KEGG pathways in GSEA and ORA was considered with an adjusted *p*<0.01.

DEG downregulated in INS+CD3-sAAb+ islets vs. ND were also decreased compared to the T1D group and included secreted factors pertaining to immune responses and response to biotic stimulus, including genes encoding complement proteins and chemokines (**Figure 3C**). There was also downregulation of the IL-1 receptor *IL1R1* as well as sphingosine-1-phosphate receptors *S1PR1* and *S1PR3*, suggesting negative regulation of immune cell trafficking. Radial plots enable visualization of average islet gene expression patterns across the four groups analyzed (i.e., ND, sAAb, mAAb, T1D) (**Figure 3D**). The downregulation of genes encoding complement proteins was the greatest in sAAb+ vs. ND islets. A progressive recovery in some complement protein gene expression was observed in the pseudo-time stages from sAAb+ to T1D, where T1D and ND islet gene expression levels were comparable. This may result from a reduction in β-cells in INS+ islets as observed by insulin stain area (**Supplemental Figure 3C**).

We also observed downregulation of genes involved in ECM remodeling and cell adhesion (**Figure 3C**). Specifically, sAAb+ islets downregulated genes encoding proteins important in cell attachment and motility (integrins, galectins, fibronectin, osteopontin), ECM structure (laminins, collagens), and tissue remodeling (matrix metalloproteases (MMPs): *MMP2, MMP7, MMP14*). Members of the REG family, including *REG1A, REG1B,* and *REG3G,* as well as ADAM proteins *ADAMTS1* and *ADAMTS9,* were also reduced (**Figure 3C**). These genes followed a similar trend as complement proteins: their expression was downregulated in sAAb+ islets and progressively increased in mAAb+ and T1D islets (**Figure 3D**).

sAAb+ islets also downregulated regulators of inflammation as well as signal transducing molecules. The tumor necrosis factor (TNF) receptor superfamily members *TNFRSF1A* (TNFR1) and *TNFRSF21* (DR6), which mediate responses to TNFα and related ligands, were consistently downregulated across all three clinical phenotype (i.e., sAAb+, mAAb+, T1D) vs. ND (**Figure 3D**). A similar pattern was observed for the NF-κB signaling regulator CHUK, the kinase of the canonical NF-κB pathway, but not of the transcription factor *NFIB* nor the negative regulator *TNFAIP3* (A20). *NFIB* and *TNFAIP3* gradually increased in expression to comparable (*TNFAIP3*) or higher levels (*NFIB*) in T1D than ND islets (**Figure 3D**). JAK/STAT signaling through STAT3 was also downregulated in sAAb+ islets, along with the alpha and beta subunits for both the type I and type II interferon receptors (*IFNAR1, IFNAR2, IFNGR1, IFNGR2*) (**Figure 3C-D**). Their expression gradually increased from sAAb+ to T1D but did not reach ND levels at the T1D stage.

### Protein homeostasis is disrupted in sAAb+ and mAAb+ islets, including suppression of translation as well as decreased ER stress and UPR genes

Significant similarities existed comparing DEG from ND to sAAb+ or mAAb+ individuals. We performed GSEA with identical DEG from the sAAb+ vs. ND and mAAb+ vs. ND gene sets (2,207 DEG; **Figure 2C**).

Among the top significantly enriched BP were the regulation of cell adhesion, catalytic activity, protein metabolic processes, as well as those noted for suppression of angiogenesis and proteolysis (**Supplemental Figure 8A**, **Supplemental Table 17**). Notably, the top three significantly enriched and suppressed KEGG pathways were metabolic, protein processing in the ER, and PI3K-Akt signaling pathways (**Supplemental Figure 8B**, **Supplemental Table 18**). Reduced protein processing in the ER was among the top five DE pathways shared between sAAb+, mAAb+, and T1D (**Figure 3C**). **Supplemental Figure 8C** and **Supplemental Table 19** present the KEGG pathway ORA for shared DEG between sAAb+ and mAAb+ comparisons.

BP related to protein processing in the ER were also downregulated in AAb+ groups (**Supplemental Figure 8D**, **Supplemental Table 20**). Protein localization to ER, ER to Golgi vesicle-mediated transport, positive regulation of translation, translational initiation, and lysosome organization were all reduced in sAAb+ and mAAb+ vs. ND groups (**Figure 4A**: heatmap of representative DEG in the BP; **Supplemental Figure 9A**: all genes). These data are consistent with our recent publications using global proteomics as well as targeted immunofluorescence (11,28). The enriched DEG were downregulated and included genes comprising four eukaryotic translation initiation factors (eIF): eIF2, eIF3, eIF4, and eIF6. Similarly, regulators of eIF function and/or translation were reduced, including the kinase *EIF2AK1*, the binding protein *EIF4EBP2,* argonaute 2 (*AGO2*), the master regulator *MTOR,* and *DDX3X*. Accordingly, downregulation of genes encoding the eukaryotic ribosome, including members of the 40S and 60S ribosomal subunits, was observed. These data confirm results in our recent islet proteomic profile involving LMD followed by LC-MS from three mAAb+ nPOD cases, including two cases analyzed here (cases 6080, 6158) (28).

**Figure 4.**
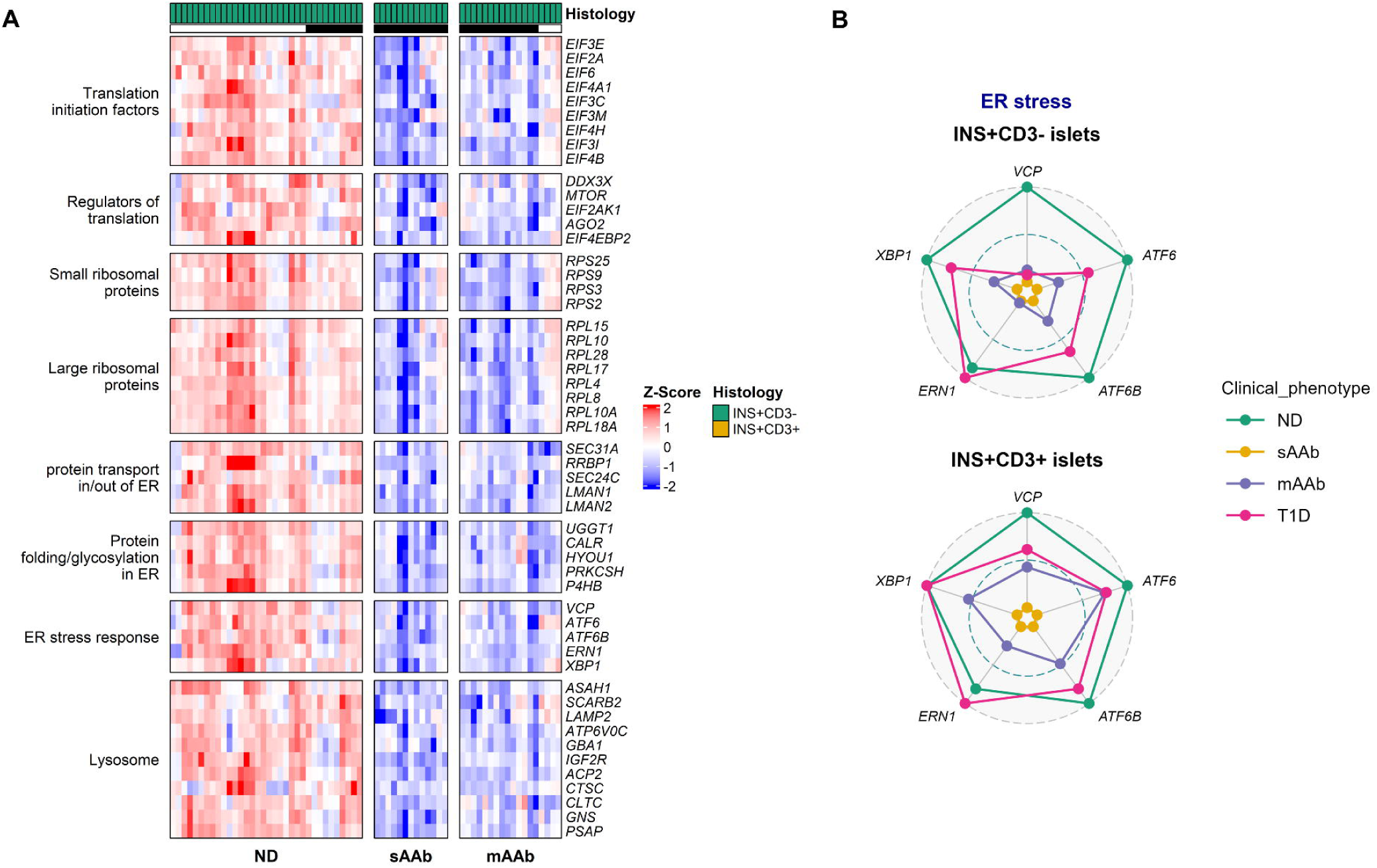
INS+CD3-islets from both sAAb+ and mAAb+ cases display downregulated translation and protein processing in the ER. **(A)** Heatmap of representative, enriched DEGs of dysregulated pathways in both INS+CD3-sAAb+ and mAAb+ islets. Gene expression in the heatmap is shown as z-scores. Black line indicates islets from age-matched subcohort. **(B)** Radial plots of islet average gene expression across clinical groups for enriched ER stress genes in INS+CD3-(top) and INS+CD3+ (bottom) islets. Gene expression in radial plots is scaled from 0 to 1, where 0 and 1 represent the minimum and maximum average gene expression values, respectively.

Consistent with translational repression, both sAAb+ and mAAb+ islets presented downregulation of protein localization to ER and protein processing in the ER (**Supplemental Figure 8C-D**). These pathways included genes involved in ER transport, folding, glycosylation, transport to Golgi, ER-associated degradation, and ER stress response genes (*ATF6, ATF6B, ERN1, XBP1*), among others (**Figure 4A**, **Supplemental Figure 9B**). These data confirm our global proteomics results and immunostaining that demonstrated decreased expression of ER-associated proteins, including ER stress (11,28). The lysosomal KEGG pathway was also suppressed in sAAb+ and mAAb+ islets (**Figure 4A**, **Supplemental Figure 10**). This pathway included the downregulation of lysosomal membrane proteins, lysosomal enzymes (proteases, glycosidases, sulfatase, lipases, and others), and enzyme trafficking genes.

Since we observed the significant downregulation of ER stress-related genes in sAAb+ and mAAb+ INS+CD3-islets, we then explored their gene expression in INS+CD3+ (mAAb+ and T1D) islets. ER stress in infiltrated islets was comparable to that of non-infiltrated islets and did not increase in T1D islets post-infiltration (**Figure 4B**). Correlation analysis showed a significant, moderate positive correlation of ER stress markers *XBP1* and *ERN1* with *CD68* (macrophages) but not *CD3E* (T cells) in INS+CD3-islets (**Supplemental Figure 11A**). Interestingly, after T-cell infiltration (INS+CD3+ islets, **Supplemental Figure 11B**), the positive correlation was only significant for *XBP1* with both *CD68* and *CD3E* expression. These results suggest that the expression levels of ER stress markers may be influenced by interactions with macrophages or innate inflammation, as they gradually increase from sAAb+ to T1D islets, but reach similar levels in T1D as ND islets (**Figure 4B**). This corroborates *in vitro* studies, where ER stress markers are upregulated after pro-inflammatory cytokine treatment of both human β-cells or isolated islets (29,30). Nonetheless, our results and others (11,28,31,32) suggest the inflammatory milieu *in situ* does not indicate that ER stress is elevated and contributing to T1D pathogenesis.

### Mitochondrial metabolism is downregulated in mAAb+ islets compared to ND persons

The β-cell functional deficit in mAAb+ individuals who progress to T1D is inversely correlated with the number of AAb, with a steeper decline in FPIR being associated with T1D gene risk loci outside the class II HLA region (8,9). Hence, we investigated the transcriptional changes uniquely DE comparing mAAb+ and ND islets (1,080 DEGs; **Figure 2C**). GSEA led to similar BP as described above, including cellular location, transport, and chromosome organization (**Supplemental Figure 12A**, **Supplemental Table 21**). Likewise, KEGG pathway ORA showed the enrichment of the ribosome pathway, viral responses, and autoimmune diseases (**Supplemental Figure 12B**, **Supplemental Table 22**). The genes associated with these pathways corresponded to other small and large ribosomal proteins and several histone genes, including histone core H2A, H2B, H3, and H4 families as well as histone variant genes (**Supplemental Figure 12C**).

While the suppression of protein translation and processing was prominent in sAAb+ and mAAb+ islets, the term “metabolic pathways” was also among the most significantly over-represented and suppressed pathways (**Supplemental Figure 5B-C**). Indeed, the BP “generation of precursor metabolites and energy” was enriched in mAAb+ vs. ND islets (**Supplemental Figure 5C**). Under this BP, we observed the significant suppression of genes involved in glycolysis (*PGM1/2, GAPDH, GPI, PKM*), mitochondrial metabolism and/or TCA cycle (*CS, IDH1/2, IDH3A/B/G, OGDH, ME2, SLC25A3/6/10/23/24/39/40, ACO2*), electron transport chain (ETC) complex I (*NDUFA9, NDUFA11, NDUFB7, NDUFV1*), complex II (*SDHAF2*, *SDHAF3*), complex III (*CYCY*), complex IV (*COX10, COX15*), and complex 5 (*ATP5F1A, ATP5F1B*) (**Figure 5A**).

**Figure 5.**
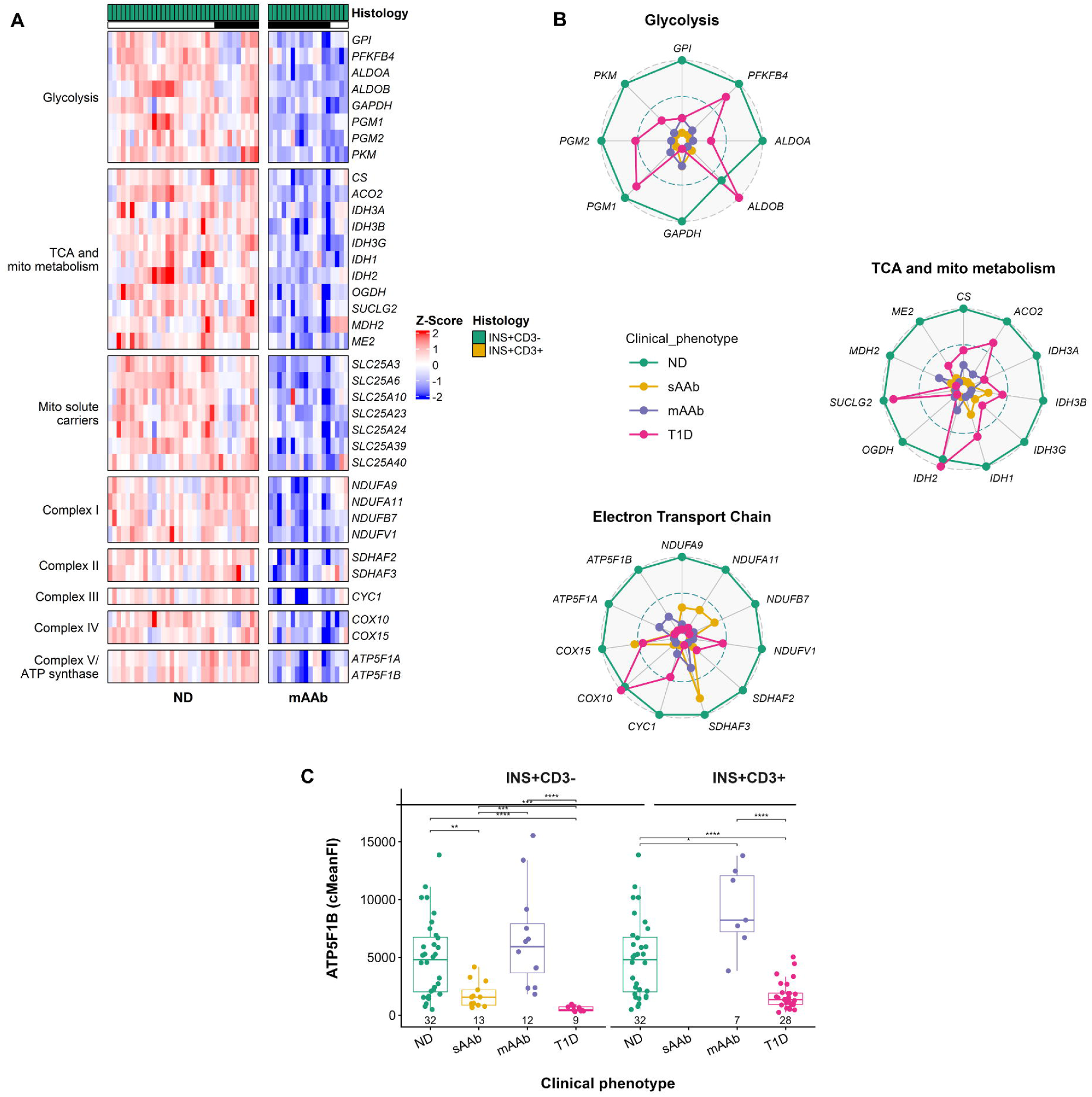
Metabolic pathways are downregulated in INS+CD3-islets from mAAb cases. **(A)** Heatmap of representative, enriched DEGs of dysregulated pathways in INS+CD3-mAAb islets. Gene expression in the heatmap is shown as z-scores. Black line indicates islets from age-matched subcohort. **(B)** Radial plots of islet average gene expression across clinical groups for enriched DEGs from A. Gene expression in radial plots is scaled from 0 to 1, where 0 and 1 represent the minimum and maximum average gene expression values, respectively. **(C)** Quantification of islet ATP5F1B by immunofluorescence staining of human pancreas sections in INS+CD3-(left) and INS+CD3+ (right) islets. For comparisons ND INS+CD3-appears in both left and right panels. The immunofluorescence signal was corrected by subtracting the acinar tissue signal from the islet region (cMeanFI). Statistical significance of cMeanFI was assessed by using the non-parametric Wilcoxon test. Multiple testing correction was performed with the Benjamini-Hochberg (BH) method. Adjusted *p*: ∗*p*<0.05, ∗∗*p*<0.01, ∗∗∗*p*<0.001, ∗∗∗∗*p*<0.0001.

The suppression of mitochondrial bioenergetics-related genes was present in both sAAb+ and mAAb+ persons (**Figure 5B**); however, mAAb+ islets demonstrated more pronounced dysregulated gene expression (**Figure 6**). Specifically, mAAb+ islets presented greater downregulation of genes comprising the ETC complex I and II. These results suggest transcriptional disruptions in glucose metabolism and/or mitochondrial bioenergetics may start at the sAAb+ stage and increase in magnitude or become combinatorial in mAAb+ individuals, as reported by clinical studies (3–7). To determine if the gene expression of these metabolic-centric genes changes post-infiltration, we evaluated the transcriptional levels of rate-limiting enzyme-encoding genes (**Figure 5A**). Overall, we observed similar gene expression magnitudes for *PKM, CS, IDH3A, IDH3B, IDH3G, OGDH,* and *ATP5F1B* in islets irrespective of T-cell infiltration (**Supplemental Figure 13A-B**).

**Figure 6.**
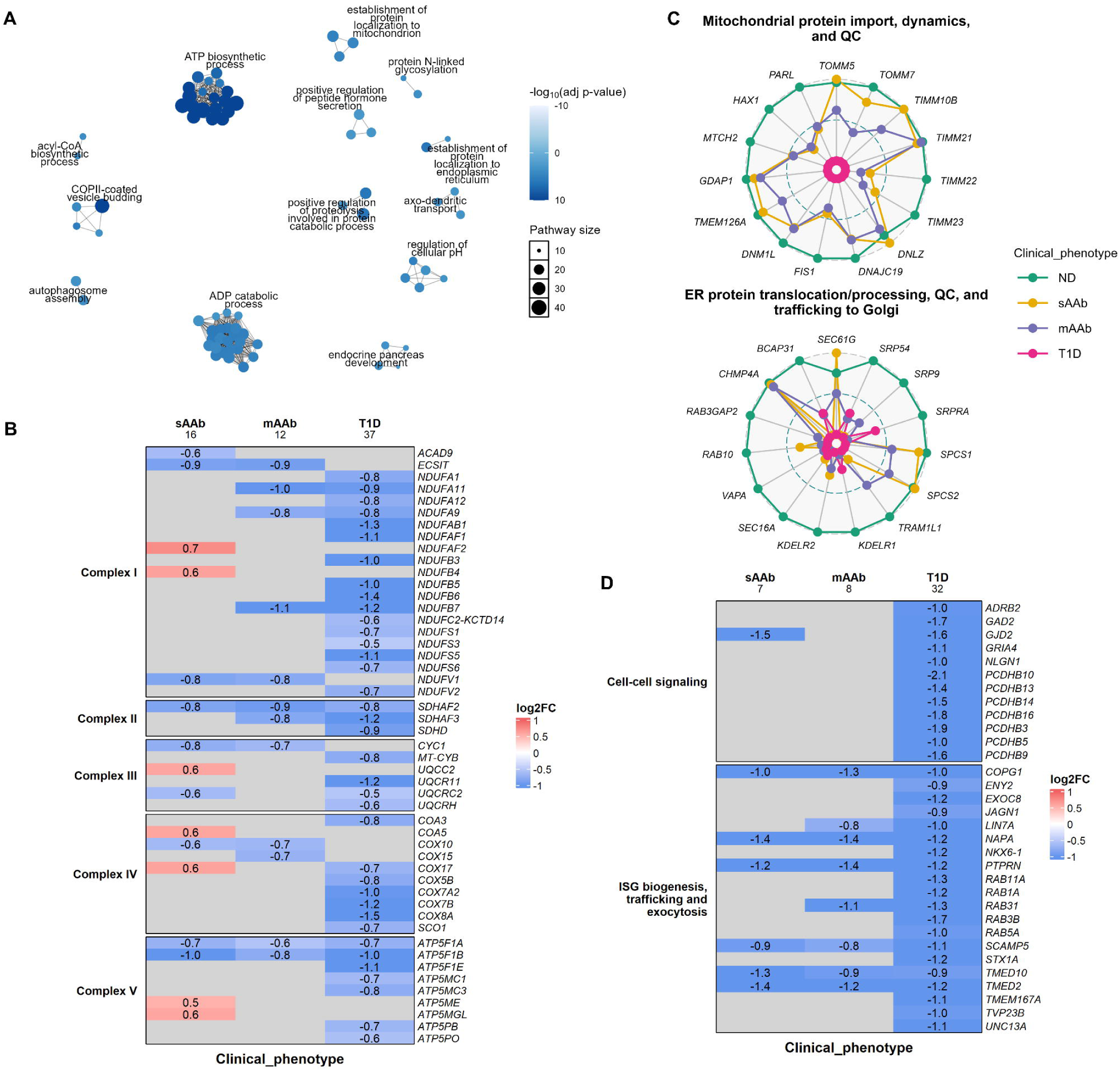
INS+CD3-islets from T1D cases downregulate mitochondrial bioenergetics and hormone secretory pathways. **(A)** Pathway clustering of the top 100 over-represented biological processes (BP) among all DEGs in the INS+CD3-T1D vs. ND comparison. **(B)** Heatmap of differential expression of electron transport chain (ETC) complex I-V genes across sAAb, mAAb and T1D islets. **(C)** Radial plots of scaled (0–1), islet average gene expression across clinical groups for additional enriched DEGs in the INS+CD3-T1D vs. ND comparison. Gene expression in radial plots is scaled from 0 to 1, where 0 and 1 represent the minimum and maximum average gene expression values, respectively. **(D)** Heatmap of differential expression of hormone secretion pathways across single(s), multiple(m) AAb and T1D islets. Gene expression in the heatmaps is shown as log2FC. The number below each clinical group label in the heatmaps represents the total number of DEGs per group.

Our unique islet phenotyping strategy allowed us to stain for ATP5F1B protein expression in subsequent serial sections to those used for single islet transcriptomics. We observed a similar pattern of expression as the transcriptomics data; namely. ATP5F1B protein expression significantly decreased in sAAb+ vs. ND islets, significantly increased from sAAb+ to mAAb+ islets, and was significantly decreased from mAAb+ in comparison to T1D islets (**Figure 5C**). Within clinical groups, ATP5F1B protein levels were identical in non-infiltrated and infiltrated islets (**Figure 5C**).

### Mitochondrial homeostasis is disrupted in T1D islets irrespective of T-cell infiltration

The loss of FPIR in recent-onset T1D islets occurs similarly in non-infiltrated and current T-cell-infiltrated islets. (11). To determine gene expression changes underlying loss of FPIR, we first performed GSEA and ORA with those genes uniquely DE in T1D INS+CD3-islets vs. ND (861 DEGs; **Figure 2C**). We observed suppression of cell-cell signaling (**Supplemental Figure 14A**, **Supplemental Table 23**) as well as reduced transcripts for genes involved in both ETC and OXPHOS (**Supplemental Figure 14B**, **Supplemental Table 24**). Neurodegeneration pathways were likely enriched with KEGG pathways due to the high content of ETC and OXPHOS transcripts in those BP (**Supplemental Figure 14C**, **Supplemental Table 25**) as reported in whole islet RNAseq (33). To describe all pathways dysregulated in T1D islets, we analyzed all DEG for the T1D vs. ND comparison (1,508 DEGs; **Figure 2A**). Pathway clustering of the top 100 over-represented BPs revealed the most significant finding was ATP biosynthetic process (**Figure 6A**, **Supplemental Table 10**). Other significantly enriched processes included the regulation of hormone levels and protein localization to mitochondria and ER (**Figure 6A**, **Supplemental Figure 5E**). Accordingly, the KEGG pathways enriched for all DEG in T1D vs. ND subjects involved reductions in OXPHOS, proteasome, and protein processing in the islet ER (**Supplemental Figure 5F**, **Supplemental Table 10**).

We observed the downregulation of multiple nuclear-encoded OXPHOS genes, including members across all ETC complexes and ATP synthase, in T1D vs. ND islets (**Figure 6B**, **Supplemental Figure 15**). Those genes were noted as downregulated, in addition to the reduced expression of glycolytic and TCA-related genes (**Figure 5B**). Indeed, the number of downregulated DEG involved in ATP biogenesis was 3-4 times greater than those observed in sAAb+ and/or mAAb+ islets (**Figure 6B**). T1D islets downregulated 16 out of 49 complex I genes and the complex II genes *SDHC*, *SDHD, SDHAF2 and SDHAF3*. In concordance with the gene expression results, we previously observed decreased SDHC protein by immunofluorescence of mAAb+ and T1D islets (11). In the current study, T1D INS+CD3-islets also suggested downregulation of complex III structural subunits along with seven of 33 complex IV genes. Seven of the 19 F1F0 ATP synthase subunits were also suppressed, including *ATP5F1A*, *ATP5F1B*, *ATP5MC1, ATP5MC3*, *ATP5PB,* and *ATP5PO*. ATP5F1B was decreased in the immunostained serial sections (**Figure 5C**). Expression levels of the ETC genes remained downregulated in T1D islets post-T-cell infiltration (**Supplemental Figure 16A**). Consistent with this, the KEGG pathways suppressed in T1D INS+CD3+ vs. ND INS+CD3-included metabolic, neurodegenerative diseases, and OXPHOS (**Supplemental Figure 16C**, **Supplemental Table 27**).

In addition to the suppression of mitochondrial bioenergetics, T1D INS+CD3-islets also presented downregulated expression of genes involved in mitochondrial protein import, dynamics, and QC (**Figure 6C**). Those genes included members of the translocase of the outer and inner mitochondrial membrane (i.e., TOMM and TIMM) complexes, including the core components of TIMM22 and TIMM23, and co-chaperone *DNAJC19*. The assembly factor *TMEM126A* that aids in inner membrane protein complexes formation, as well as *DNLZ,* which regulates chaperone activity, were also downregulated. Moreover, genes involved in mitochondrial fission and QC followed a similar trend and included the fission initiator *GDAP1*, adaptor recruiter *FIS1*, GTPase *DNM1L,* and apoptosis regulators *MTCH2* and *HAX1*.

### Hormone secretory pathways are similarly suppressed in non-infiltrated and infiltrated T1D islets compared to ND

Concomitant to the overall suppression of ATP biogenesis, T1D INS+CD3-islets exhibited reduced BPs in cell-cell signaling and regulation of hormone levels and secretion (**Supplemental Figure 5E**, **Supplemental Table 9**), and pathway clustering revealed the enrichment of proteolysis, protein localization to the ER, and peptide hormone secretion (**Figure 6A**). Similar to sAAb+ and mAAb+ islets, T1D INS+CD3-islets presented dysregulated protein homeostasis in their ER. Notably, T1D islets downregulated genes involved in targeting proteins to the ER, translocation to the ER, maturation in the ER, trafficking to Golgi, and packing into secretory vesicles for exocytosis (**Figure 6C**). There was also a vast downregulation of genes tightly involved in the insulin granule biogenesis, trafficking, and exocytosis (**Figure 6D**), including COPI/COPII-mediated vesicle trafficking genes involved in vesicle sorting and mobilization, the exocyst complex member *EXOC8*, members of the exocytosis machinery, and core SNARE proteins α-SNAP (*NAPA*), Munc13-1 (*UNC13A*), and syntaxin-1A (*STX1A*). The insulin granule transmembrane protein IA-2 (*PTPRN*) was also downregulated along with the pro-insulin converting enzymes *CPE* (logFC=-0.9, adjusted *p*=0.03) and *PCSK1* (logFC=-2.1, adjusted *p*=0.0013). T1D islets downregulated genes regulating insulin secretion, including the transcription factor *NKX6-1*, nuclear factor *ENY2,* and glucagon-like peptide-1 receptor (*GLP1R*) (Figure 6D). Genes involved in islet cell-cell signaling were also down, including the glutamate decarboxylase 65 (*GAD2*), which synthesizes GABA, and the gap junction protein connexin-36 (*GJD2*) (**Figure 6D**).

The downregulation of these genes and processes did not change in T1D islets with infiltration (**Supplemental Figure 16B**). Indeed, the ORA analysis with T1D INS+CD3-islets vs. ND revealed that these BPs remained suppressed while immune-related pathways were activated (**Supplemental Figure 16C-D**).

### Pseudo-transition to T1D is associated with activation of immune and stress responses

The BP enriched in the pseudo-transition from mAAb+ to T1D included activation of proteolysis, stress and immune responses (**Supplemental Figure 5G, Supplemental Table 11, Supplemental Figure 17**). As expected, HLA Class I (*HLA-C*) was upregulated. HLA Class II (*HLA-DRA, HLA-DRB1, HLA-DMA, HLA-DMB*) genes were upregulated, indicating increased antigen presentation in T1D islets and presence of activated antigen-presenting cells (**Figure 7A-B**). As expected, the upregulation of class I and class II HLA positively correlated with *CD68* expression (surrogate for macrophages) but not with *CD3E* (T cells) in INS+CD3-islets (**Figure 7C**) (22). However, the association of *HLA-C* and *CD3E* became significant in T-cell infiltrated islets (**Figure 7C**). Consistent with the increase of antigen-presenting cells from mAAb+ to T1D islets, there was also upregulation of chemotaxis-related genes, including *CCR1* and *CXCL6.* The pseudo-transition to T1D in INS+CD3-islets also revealed upregulation of interferon-stimulated genes (ISGs) *TRIM25, SP100, DAP,* and *DAPK1,* associated with anti-viral responses and apoptosis (**Figure 7A**). Under the BP of proteolysis, we observed the significant upregulation of ECM remodeling enzymes and tissue proteases and regulators (*MMP14*, *KLK1, SPINK1, SPINT1*) (7A). We also observed upregulation of *CD44* (logFC = 2.9), the hyaluronan (HA) receptor. Here, the gene expression patterns of *CD44* and *MMP14* were similar to other ECM-related genes as described previously for sAAb+ islets (**Figure 7B**). Hence, we hypothesize that as T1D progression occurs, the peri-islet capsule becomes compromised, facilitating immune cell infiltration.

**Figure 7.**
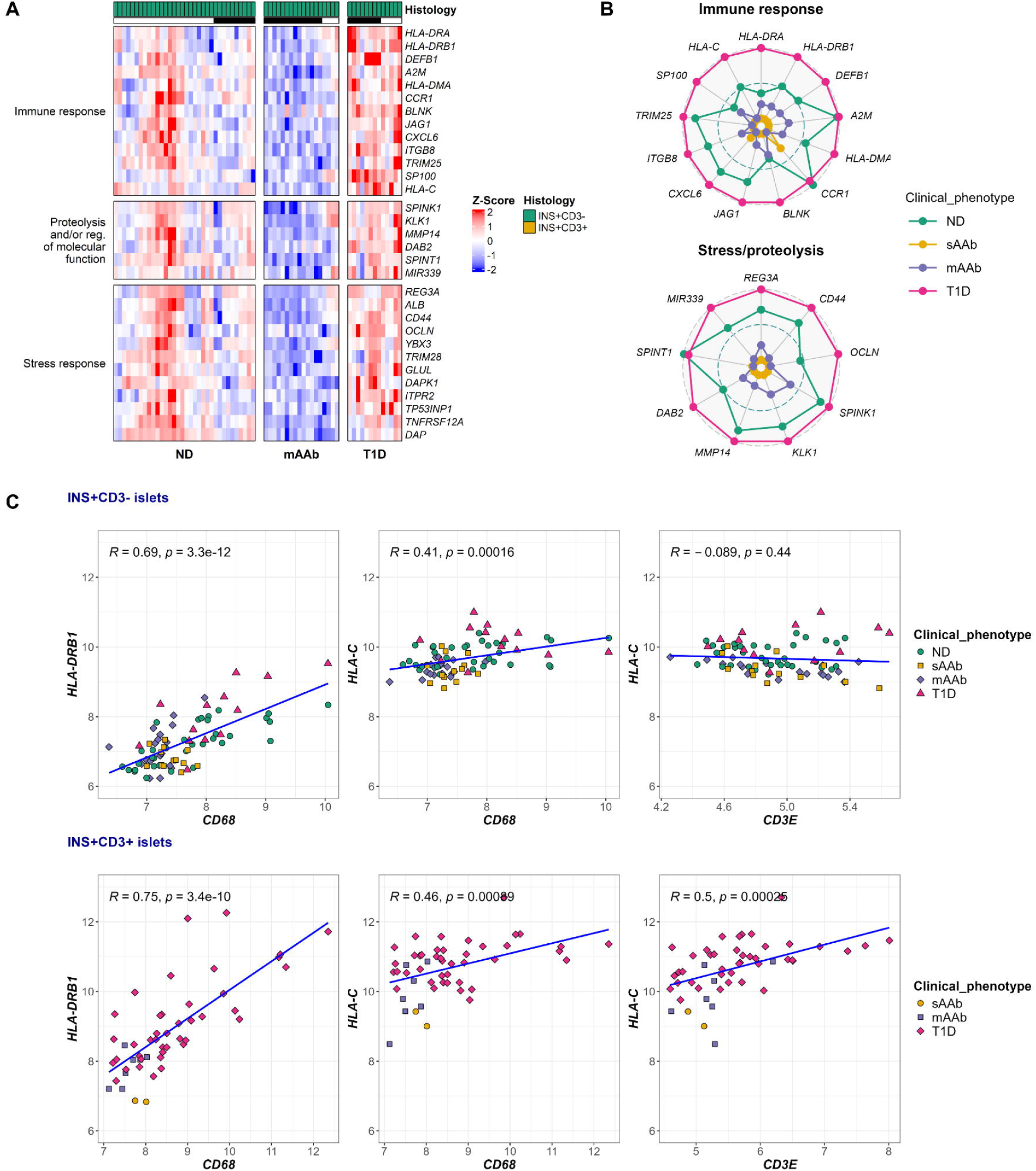
INS+CD3-islets from T1D cases activate immune and stress response. **(A)** Heatmap of representative, enriched DEGs of dysregulated pathways in INS+CD3-T1D islets when compared to mAAb islets. Gene expression in the heatmap is shown as z-scores. Black line indicates islets from age-matched subcohort. **(B)** Radial plots of scaled (0–1), islet average gene expression across clinical groups for enriched DEGs from A. Gene expression in radial plots is scaled from 0 to 1, where 0 and 1 represent the minimum and maximum average gene expression values, respectively. **(C)** Pearson correlation plots of islet gene expression between HLA class I and surrogate genes for macrophages and T-cells. Top C panel displays INS+CD3-islets, bottom C panel displays INS+CD3+ islets. Statistical significance of Pearson correlations was considered with a *p*<0.05.

### T-cell infiltration drastically increases upregulation of antigen processing and presentation genes

To understand the islet environment during immune cell infiltration, we compared the DEGs in T-cell infiltrated islets (CD3+) vs. non-infiltrated (CD3-) for the T1D and mAAb+ groups (Table S29). Under the stringent cutoffs applied, there were only two highly upregulated genes in infiltrated mAAb+ islets, both encoding G protein-coupled receptors (GPCRs): *ADGRG7* (logFC=2.8) and *OR11H1* (logFC=1.5) (**Figure 8A**). However, the mAAb+ CD3+ versus CD3− comparison included insulitic islets from only two donors. These findings may suggest altered GPCR-associated signaling in infiltrated mAAb+ islets, but the limited donor representation precludes definitive interpretation.

**Figure 8.**
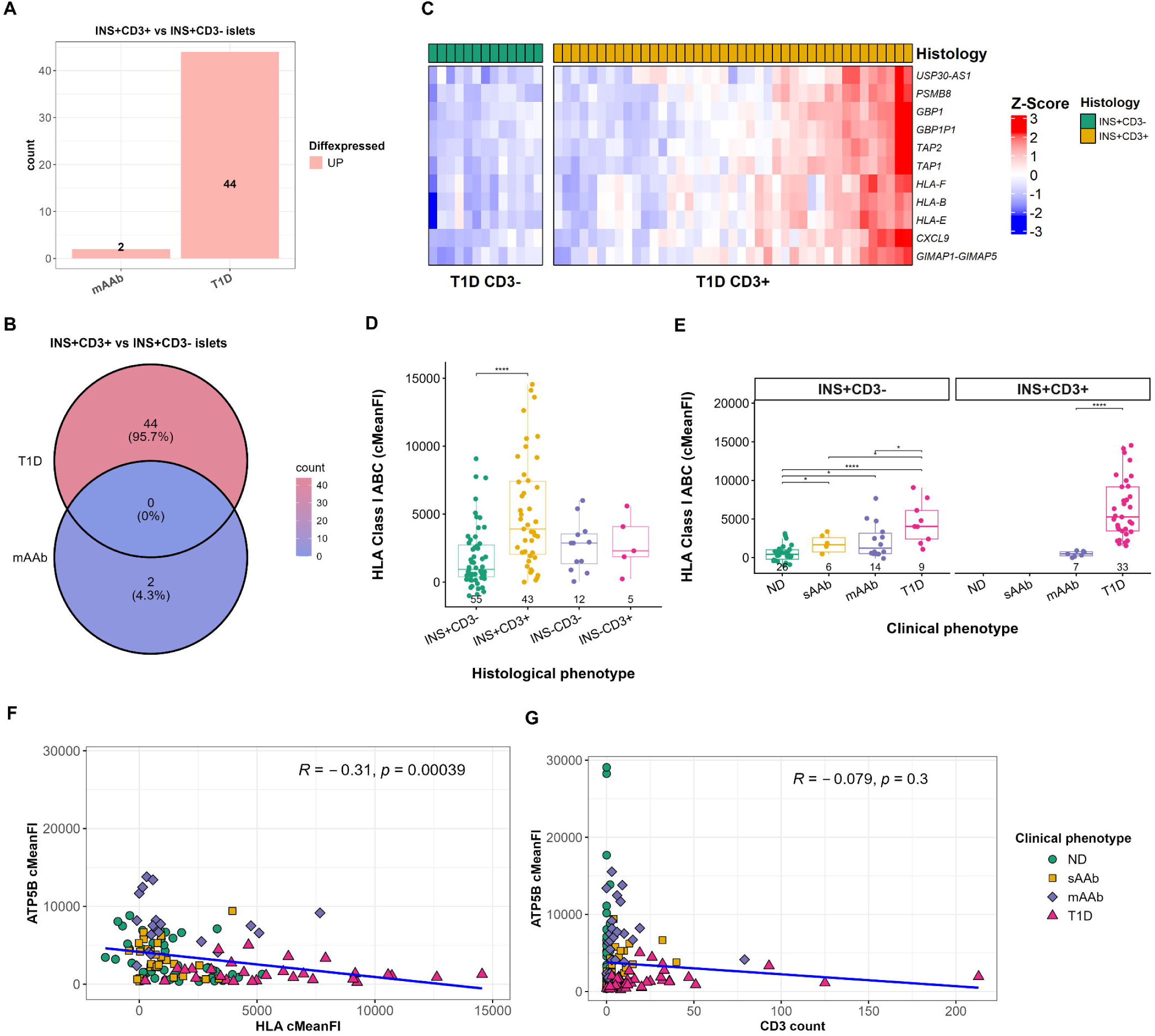
Transcriptional changes occurring post T-cell infiltration within islets from mAAb positive and T1D cases. **(A)** Number of DEGs when comparing INS+CD3+ vs. INS+CD3-islets from mAAb and T1D groups respectively. DEGs were defined as genes with an absolute logFC˃0.5 and an adjusted *p*˂0.001. **(B)** Venn diagram of DEGs from A. **(C)** Heatmap of DEGs occurring in T1D islets post-infiltration. Gene expression in the heatmap is shown as z-scores. **(D)** Quantification of islet HLA Class I ABC by immunofluorescence staining of human pancreas sections across the islet histological and **(E)** clinical phenotypes. The immunofluorescence signal was corrected by subtracting the acinar tissue signal from the islet region (cMeanFI). **(F)** Pearson correlation plot of ATP5B cMeanFI vs. HLA Class I cMeanFI and **(G)** CD3+ cell count. Statistical significance of cMeanFI was assessed by using the non-parametric Wilcoxon test. Multiple testing correction was performed with the Benjamini-Hochberg (BH) method. Adjusted *p*: ∗*p*<0.05, ∗∗*p*<0.01, ∗∗∗*p*<0.001, ∗∗∗∗*p*<0.0001. Statistical significance of Pearson correlations was considered with a *p*<0.05.

The comparisons between T1D islets that are T-cell infiltrated versus those that are not, revealed 44 DEG with an adjusted p≤0.001 (**Figure 8A**). There was no overlap in the DEG between infiltrated mAAb+ and T1D islets vs. their non-infiltrated counterparts under the defined cutoffs (**Figure 8B**). We observed the upregulation of several ISGs including Class I HLA (*HLA-B, HLA-E, HLA-F*), antigen processing and immunoproteasome (*PSMB8, TAP1, TAP2*), *CXCL9,* and interferon-inducible GTPase *GBP1* in CD3+ T1D islets (**Figure 8C**). We confirmed the significant increase in HLA-Class I protein expression by immunostaining in INS+CD3+ islets (**Figure 8D**). In addition, INS+CD3-mAAb+ and T1D islets also presented significantly higher levels of HLA-Class I when compared to ND (**Figure 8E**). This agrees with multiple previous studies (11,32,34). We also observed a significant negative correlation of Class I HLA and ATP5F1B protein expression (Figure 8F), suggesting that as mitochondrial function declines, Class I HLA expression increases. In support of this interpretation, no correlation was observed between the protein levels of ATP5F1B and T-cell count/islet (CD3+ cells) (**Figure 8G**).

### Infiltrated T1D islets upregulate immune- and anti-viral-related genes

Our analyses of INS+CD3-islets demonstrated that β-cell metabolic and secretory pathways are transcriptionally downregulated in T1D islets, both before and during infiltration. To describe DEG changes occurring in infiltrated islets, we compared mAAb+ and T1D INS+CD3+ islets to the ND INS+CD3-control islets, respectively (**Supplemental Table 29**). Only 20 genes were DE in mAAb+ INS+CD3+ vs. ND under the stringent adjusted *p* cutoff (<0.001) (**Figure 9A**). Of those, six DEG were also present in T1D INS+CD3+ islets (**Figure 9B**). Those genes changed in the same direction (downregulation) in mAAb+ and T1D INS+CD3+ islets and corresponded to genes involved in ER stress and protein processing (*SLC33A1, HSPA13*), transcriptional regulation (*CITED2, ZMAT4*), apoptosis (*RASSF6*), and mitochondrial metabolism (*BDH1*). Conversely, we observed over 2,500 DEG in T1D INS+CD3+ vs. ND INS+CD3-islets (**Figure 9A**). Of those, 928 DEG were upregulated. The top activated BP corresponded to cell killing, adaptive immune response, and regulation of immune response (**Figure 9C-D**, **Supplemental Table 30**). INS+CD3+ T1D islets presented broad activation of immune pathways, consistent with increased antigen presentation, interferon signaling, leukocyte recruitment, and cytotoxic effector function (**Figure 9D**, **Supplemental Figure 16 C-D**).

**Figure 9.**
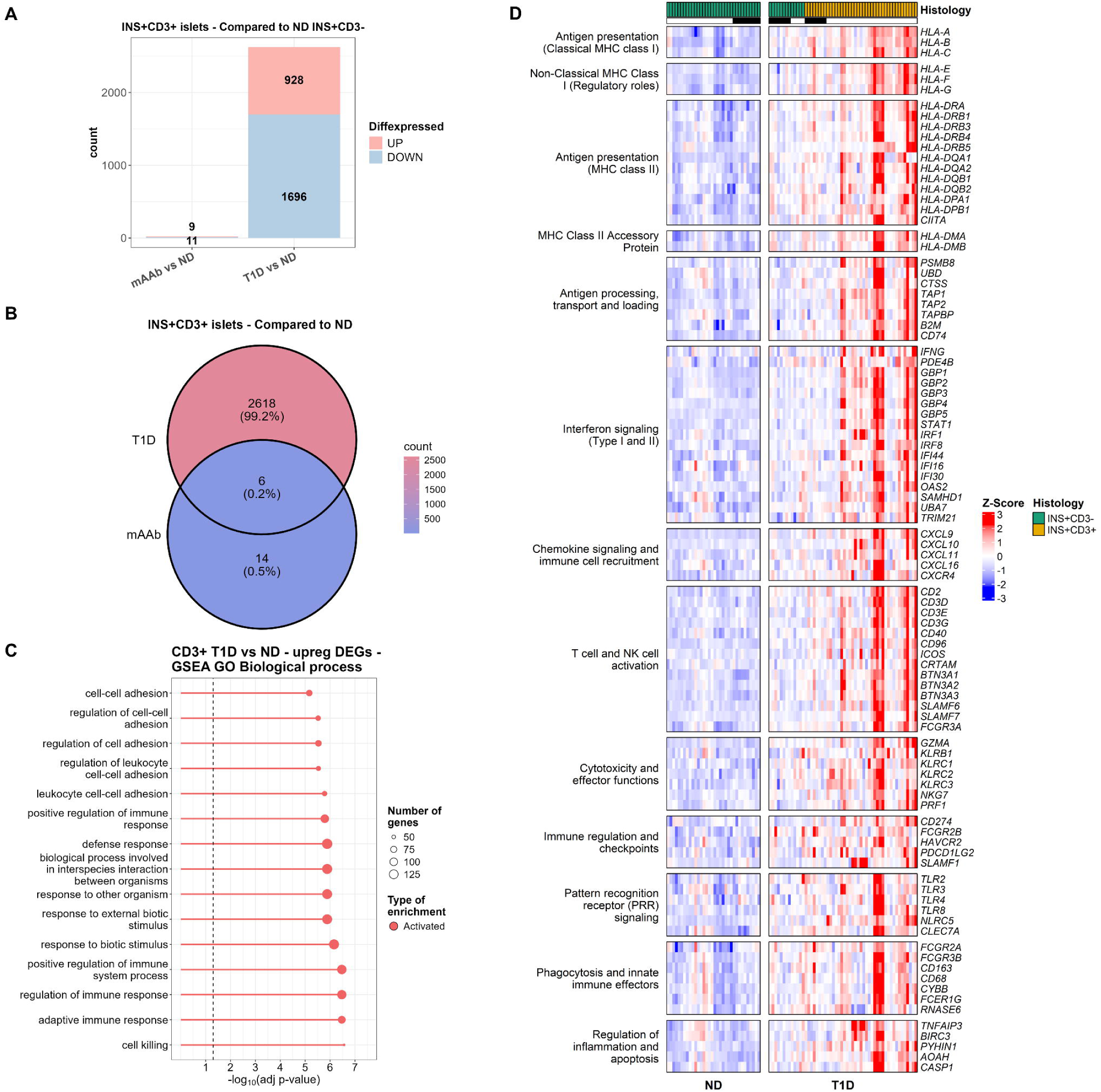
T1D islets activate multiple immune-related pathways post T-cell infiltration. **(A)** Number of DEGs in INS+CD3+ mAAb and T1D islets, respectively, when compared to INS+CD3-ND. DEGs were defined as genes with an absolute logFC˃0.5 and an adjusted *p*˂0.001. **(B)** Venn diagram of DEGs from A. **(C)** Gene set enrichment analysis (GSEA) with gene ontology (GO) biological processes (BP) of upregulated DEGs in the INS+CD3+ T1D vs. INS+CD3-ND comparison. Statistical significance for BPs was considered with an adj. p value cutoff < 0.01. **(D)** Heatmap of representative DEGs from enriched BPs and KEGG pathways in the INS+CD3+ T1D vs. INS+CD3-ND comparison. Gene expression in the heatmap is shown as z-scores. Black line indicates islets from age-matched subcohort.

T1D INS+CD3+ islets showed increased expression of classical and non-classical HLA Class I genes (*HLA-A/B/C/E/F/G*), accompanied by increased expression of genes involved in antigen processing, transport, and loading (*PSMB8, TAP1, TAP2, TAPBP, B2M,* others), as well as induction of HLA Class II genes (*HLA-DRA, HLA-DRB1,3-5, HLA-DQA1-2, HLA-DQB1-2, HLA-DPA1/DPB1*). INS+CD3+ T1D islets presented a strong interferon-driven inflammatory signature, with upregulation of *IFNG, STAT1, IRF1, IRF8, IFI44, IFI6, IFI16, OAS2, SAMHD1, TRIM21*, and multiple guanylate-binding proteins (*GBP1–5*). In parallel and consistent with the interferon-signature, there was increased expression of chemokines involved in immune-cell recruitment, including *CXCL9/10/11/16,* and the chemokine receptor *CXCR4*.

T1D INS+CD3+ islets also presented upregulation of several T-cell and NK-cell markers as well as cytotoxic/effector genes, and genes involved in regulation of inflammation and apoptosis (**Figure 9**). There was significant upregulation of myeloid innate immune cell markers and pattern recognition receptor (PRR) genes. Collectively, we observed that both mAAb+ and T1D INS+CD3+ islets present downregulation of ATP biogenesis, ER stress, and protein homeostasis when compared to ND INS+CD3-islets. In addition, T1D INS+CD3+ islets continue with this signature and add a pronounced pro-inflammatory, interferon-driven signature as well as the concomitant presence of both myeloid innate and cytotoxic effector immune cells, including NK and T-cells.

## Discussion

The loss of FPIR is a major risk factor for T1D progression. The mechanisms for failed β-cell responses to glucose have remained a major knowledge gap in the development of therapies to maintain insulin secretion. Histological examination of islets in the prediabetic period of T1D has revealed that β-cell mass remains unchanged while insulitis, if present, is rare (10,32,35,36). We have recently shown using immunostaining that the loss of FPIR in T1D islets is associated with decreased proteins in the glucose-stimulated insulin secretion (GSIS) pathway and independent of acute T-cell infiltration (11). Furthermore, we also showed that dysfunctional β-cells in stage 1 T1D are inversely associated with an islet immune response (28,37). The goal of this work was to follow up on those studies by providing an in-depth transcriptional analysis of islets throughout the natural history of T1D, including the comparative analysis of islets with and without T-cell infiltration.

The advancement of multi-omics technologies over the past decade has increasingly aided in our understanding of disease pathogenesis, particularly in the heterogeneity of islet distribution and loss (10,22,32,35,38–41). However, single-cell resolution technologies pose limitations due to the inherent stress associated with islet isolation and dispersion processes from one side, and the limited number of detectably expressed genes on the other side (14,15). In addition, islets that are not intact due to immune cell infiltration and killing are not prioritized in islet isolation due to the inherent selectivity of this process. To circumvent that, we utilized a single-islet strategy using immunostaining-guided LMD to obtain transcriptomes from islets with defined phenotypes, including T-cell infiltration status. Furthermore, while spatial transcriptomics conserves the tissue architecture, a single cross section of an islet may not be fully representative of its infiltration environment. Hence, we captured whole-islet context through serial sectioning for IF and LMD (**Supplemental Figure 1**). Indeed, our approach revealed high consistency in islet phenotypes and their gene expression (**Figure 1**).

Our DE analyses in non-infiltrated islets with remaining β-cells (INS+CD3-) across the natural history of T1D revealed transcriptomic changes starting at the sAAb+ state. Strikingly, an erosion of metabolic gene expression occurred from sAAb+ cases to mAAb+, and worsened dramatically in the T1D state. Indeed, both sAAb+ and mAAb+ islets presented downregulated genes related to glycolysis and TCA (**Figure 5**), which further expanded to the ETC complexes in T1D (**Figures 5 and 7**). Those gene expression changes occurred similarly in non-infiltrated and infiltrated islets, suggesting that the progressive decline in β-cell function is uncoupled from active immune infiltration. As GSIS requires metabolism of glucose to ATP within the mitochondria, our transcriptomic analyses suggest that the loss of FPIR in T1D is associated with suppression of ATP biogenesis pathways, including glycolysis, TCA, and OXPHOS (**Figures 5 and 7**). These results are in alignment with previous single-cell RNA-seq analysis of HPAP islets (42), our recent findings by IF of nPOD pancreas tissues (11), and loss of ATP5F1B (**Figure 5C**). Notably, these results further corroborate the dysfunctional state observed in T1D islets within pancreas tissue slices (11,12). In addition, we observed decreased gene expression of core proteins belonging to the TIMM and TOMM complexes. TIMM23 depends on both the mitochondrial membrane potential and the presence of ATP to carry out its function (43). TIMM23 downregulation could be linked to the reduced OXPHOS state observed in T1D islets. Indeed, complex I inhibition has been associated with the downregulation of both TOMM20 and TIMM23 in a dopaminergic neuroblastoma cell line (44). Additionally, the downregulation of *FIS1* in the INS-1 pancreatic β-cell line resulted in reduced maximal respiratory chain capacity, production of reactive oxygen species (ROS), and the accumulation of oxidized mitochondrial proteins leading to decreased insulin secretion (45). Our results herein suggest that the marked disruption of mitochondrial homeostasis in T1D islets contributes to the overall glucose-unresponsive state historically observed in the clinic (3–9,46) and *in-situ* with pancreas tissue slices (11,12).

Concomitant with the suppression of mitochondrial bioenergetics in non-infiltrated T1D islets was the downregulation of hormone secretory pathways, including insulin granule dynamics and cell connectivity-related genes. These genes were drastically downregulated only in T1D islets. Interestingly, a study investigating soluble protein secretion in mammalian cells revealed that knocking out any member of the exocyst complex (including *EXOC8*) disrupted delivery to plasma membrane resulting in intracellular cargo accumulation in specialized secretory cells like adipocytes (47). Further, deficiency of the SNARE complex proteins Munc13-1 and syntaxin-1A has been shown to significantly reduce insulin secretion and the replenishment of insulin granules (48–50). Similar effects were observed by knocking down IA-2 (*PTPRN*) due to decreased number of insulin granules (51,52). Our findings suggest β-cells in T1D are not only metabolically impaired but also exhibit dysfunction in insulin secretory granule maturation, the exocytotic machinery, and cell connectivity. These processes are intrinsically linked, as the rise in intracellular ATP following glucose metabolism is required for the closure of ATP-sensitive K⁺ channels, membrane depolarization, Ca²⁺ influx, and subsequent insulin secretion. Thus, the loss of GSIS in T1D islets may result from the combinatorial effect of dysregulated mitochondrial bioenergetics, β-cell secretory machinery, and cell connectivity. We demonstrate that these processes are similarly impaired in infiltrated T1D islets, suggesting that those defects developed separately.

Our transcriptomic studies delineated pathways that are disrupted in the pseudo-progression of T1D. Indeed, at the sAAb+ state, we observed the downregulation of complement-, ECM remodeling-, and adhesion-related genes. Interestingly, we observed a pattern of increasing expression of these genes from sAAb+ to mAAb+ to T1D. A similar pattern was found for complement proteins in plasma of individuals before and after seroconversion, where complement proteins were higher before seroconversion but significantly decreased after the development of AAb (53,54). The downregulation of ECM remodeling genes, including MMPs in both sAAb+ and mAAb+, is consistent with recent reports of single-islet proteomics and single-cell RNA-seq analysis (37,42). Interestingly, ECM proteins, including collagen molecules, were also reduced in the plasma of individuals after seroconversion (53). Our results and others suggest that AAb+ islets present dysregulated, suppressed remodeling of ECM, potentially within the peri-islet capsule. It is tempting to speculate that the downregulation at the AAb+ stage is protective and prevents immune cell trafficking/infiltration into the islets. Nonetheless, as pseudo-transition towards T1D development occurs, this suppressive mechanism is lost. The expression of these genes increased from the sAAb+ to the T1D stage. Supporting this notion, a previous study showed that ECM components in the peri-islet capsule are lost at sites of autoimmune infiltration in the NOD mouse and human islets (55). Indeed, infiltrated islets from NOD pancreas upregulated MMPs, including *MMP2*, *MMP7,* and *MMP14* (55). Interestingly, the HA receptor *CD44* was upregulated from mAAb+ to T1D islets. CD44 is involved in stress responses and immune cell recruitment (56), and has been associated with the shift between OXPHOS and anaerobic glycolysis in cancer cells (57). Islet HA deposits have been previously reported in pancreatic tissue from sAAb+ and mAAb+ donors, and the amount of HA observed was positively correlated with the number of AAbs and independent of islet insulitis (58). Together, these findings suggest that the increased *CD44* expression from mAAb+ to T1D islets may represent a ‘cry-for-help’ in response to the dysfunctional state. However, such a response may be double-edged, as it could also facilitate immune-cell recruitment or retention within the islet microenvironment.

We also observed the persistent downregulation of translation, protein processing in the ER, and ER stress at the AAb state of T1D pathogenesis. T1D islets also presented dysregulated protein homeostasis related to ER localization, ER-to-Golgi transport, and secretory vesicle packing and trafficking, consistent with the suppressed hormone secretory processes described above. The suppression of both translation and ER-related processes in mAAb+ islets was also observed by mass spectrometry (MS)-based proteomics (28), in T1D islets by single-cell RNAseq (42), and immunostaining (11,32). Indeed, unchanged protein levels of ER stress markers have been observed in two independent studies of FFPE pancreas sections from sAAb+, mAAb+, and recent-onset T1D cases by IF (PERK and ERN1) (11) and by imaging mass cytometry (XBP1, ERN1, WFS1) (32). This is further in accord with a study using prediabetic NOD mouse pancreas tissue (31). In summary, results across several studies by multiple groups suggest that the suppressed metabolic and secretory state of T1D islets may be associated with reduced translation and protein processing in the ER, but not with increased ER stress.

In contrast to the suppressed protein homeostasis at the sAAb+ and mAAb+ states, post-transcriptional gene regulation by miRNAs and the spliceosome was activated. Remarkably, our miRNA-target interaction analysis revealed a plausible mechanism underlying the downregulated genes, as 20-50% of the miRNA targets corresponded to DE genes in sAAb+ and mAAb+ islets, which were over-represented in processes related to translation, ER stress, and cell adhesion.

Post-transcriptional gene regulation by miRNAs is critical to β-cell function, as its disruption through β-cell-specific DICER knockout results in overt diabetes (59–61). Likewise, altered alternative splicing has been negatively correlated with β-cell identity in diabetes (37,62), as well as with altered expression of T1D and T2D candidate genes (63). Hence, we hypothesize that activation of miRNA-mediated gene silencing at the AAb+ state may represent a protective mechanism that suppresses adhesion-, translation-, and ER-processing-related genes. Thus, our findings and others’ (11,28,32,42) suggest that decreased translation and ER stress are features of human T1D pathogenesis.

Upon comparing infiltrated versus non-infiltrated islets from T1D donor groups, our observations confirm the presence of an interferon-related signature in infiltrated T1D islets, including upregulation of immunoproteasome, proteins for antigen presentation, and chemokines such as *CXCL9* (11,32,34,64). Furthermore, the comparison of infiltrated T1D islets versus non-infiltrated ND revealed vast activation of immune-related pathways, along with maintaining the reductions in genes for glucose metabolism essential for FPIR, translation, and processing of proteins in the ER. Interestingly, we observed the presence of not only cytotoxic effector cells and molecules (NK and T-cells) but also markers of innate immune cells and phagocytosis, indicative of active antigen uptake, processing, and presentation. Infiltrated T1D islets also upregulated regulators of inflammation and apoptosis, including *TNFAIP3* and *CASP1*. Collectively, our results demonstrate that INS+CD3+ T1D islets exhibit an inflammatory state characterized by simultaneous induction of proinflammatory, stress response, and compensatory protective mechanisms, highlighting the complexity of the autoimmune process.

While our study presents an in-depth transcriptomic analysis of islets throughout T1D progression, several limitations exist. First, due to the rigorous QC performed, only 129 of 260 islets were analyzed. Second, our single-islet approach does not discriminate between the heterogeneity of β-cells and other cell types. Lastly, our pseudo-time comparisons represent a snapshot in the natural history of T1D. Nonetheless, our findings suggest that pancreatic β-cells in T1D progression not only become metabolically impaired but also exhibit dysfunctional mitochondrial and protein homeostasis. The transcriptional similarities between islets from sAAb+ and mAAb+, as well as T1D islets pre- and post-infiltration, provide insights into β-cell dysfunction that precede the autoimmune attack. Furthermore, we provide an extensive atlas of novel targets disrupted during T1D pathogenesis that could be exploited in future therapeutic strategies aimed at preserving or restoring β-cell function.

## Methods

### Sex as a biological variable

Our study included similar proportions of male and female pancreas organ donors, and similar findings were observed across both sexes within groups.

### Cases and pancreas tissue

Fresh frozen pancreas sections embedded in optical cutting media (OCT) were obtained from nPOD following processing by standard operating procedures in full compliance with federal and University of Florida Institutional Review Board (UF IRB) regulations for research sample use from organ donors (13). Donor groups were as follows: donors without diabetes and autoantibody negative (ND, n=16), single autoantibody positive (sAAb+, n=6), multiple autoantibody positive (mAAb+, n=5), and type 1 diabetes (T1D, n=16) (**Supplemental Table 1**). Donors were matched for age, sex, and race/ethnicity as feasible; one mAAb+ donor (6080, 69.2 years old) had no suitable matching age ND donor. Pancreas blocks were selected from those regions previously identified by immunohistochemistry to contain islets with β-cells and/or insulitis in the AAb+ and T1D donors using methods previously described (10). Insulitic islets were defined as ≥6 CD3+ cells touching the islet endocrine cells (10). A graphical representation of the methods applied to the frozen sections, as described below, is provided (**Supplemental Figure 1**).

#### Multiplex immunofluorescence (mIF)

Blocks were serially cut into either thick (10 μm) (3–4) sections, placed onto PEN-slides (Leica), or 3-4 thin (4 μm) sections placed onto Superfrost slides (Thermofisher) (Suppl. Figure 1A). Thin pancreas sections immediately before and after the thick section set were immunostained for insulin (INS), glucagon (GCG), CD3, HLA-ABC, and/or ATP5F1B and nuclei counterstained with DAPI as previously described (10,11,65). The INS, GCG, and CD3 stains permitted identification of islets within four subtypes (i.e., INS+CD3-, INS+CD3+, INS-CD3+, and INS-CD3-) based on residual β-cells and T-cell infiltration. Stained sections were scanned using a Zeiss710 laser scanning confocal microscope (Carl Zeiss). Slide section maps were generated to select islets of interest for LMD in the intervening thick sections. Islet insulin immunopositivity (INS+) included those with ≥1 β-cell in either thin section. Islet CD3+ cell counts were determined manually by 2 observers (EAB, MCT) from both sections, and the higher CD3+ islet cell count used to define CD3+ infiltration status (≥6 CD3+ cells/islet indicating insulitis or infiltration). Islet insulin, HLA, and ATP5F1B mean fluorescence intensity levels (MeanFI) were determined using FIJI image analysis software as previously described (66). Islet MeanFI were corrected by subtracting the acinar MeanFI values (cMeanFI). IF data was analyzed with the statistical tools of rstatix package v0.7.2 (https://rpkgs.datanovia.com/rstatix/) and visualized in RStudio v2024.04.0 with the graphical tools of ggpubr v0.6.0 (https://rpkgs.datanovia.com/ggpubr/), ggplot2 v3.5.1 (67), and patchwork v1.3.0 (https://patchwork.data-imaginist.com/).

### Laser microdissection and RNA isolation

Single islet LMD was performed using a Leica 7000 LMD microscope. Thick sections were removed from -80°C storage and processed for LMD as described (33). Islet autofluorescence and anatomical landmarks from section maps were used to identify single islets of interest. LMD yielded 3-4 sections per islet that were captured into 0.6 mL RNAse-free tubes. Sets of 7-25 islets per donor were microdissected, including all 4 islet subtypes as feasible. Total islet RNA was extracted using the PicoPure RNA kit (ThermoFisher). DNAse treatment was performed using recombinant DNase I (RNase-free DNAse kit, Worthington Biochemical Corporation). An aliquot (4 μL) was used for RNA QC assays (University of Florida ICBR Gene Expression and Genotyping Core Core Facility, RRID:SCR_019145)RNA concentration and quality (RIN) were assessed using an Agilent 2100 Bioanalyzer (Agilent Technologies). The remaining sample (8-12 μL) was stored at -80°C and batch shipped on dry ice to the University of Tennessee Health Science Center (Memphis, TN). A total of 260 islets were obtained across the four donor groups (ND=73, sAAb+=40, mAAb+=37, T1D=110) (**Supplemental Table 2**).

### Transcriptomic analysis

Microarray gene expression profiling (Affymetrix Human Gene 2.0 ST expression arrays (Thermo Fisher Scientific)) was performed at the University of Tennessee Health Science Center Molecular Resource Center (Memphis, TN). RNA was amplified using Nugen Pico WTA amplification kit, fragmented, and labeled using the Nugen Encore Biotin Module (Nugen). Amplified RNA samples were hybridized by Nugen protocols on the Affymetrix Fluidics Station 450 and scanned on a GCS3000 7G (Affymetrix). Raw signal intensity values from all arrays were normalized with the Guanine Cytosine Count Normalization (GCCN) and Signal Space Transformation (SST) before Robust Multi-array average (RMA) algorithm.

### Bioinformatics analysis

Data mining was performed in RStudio v2024.04.0. The SST-RMA-normalized microarray signals were variance-stabilized using the justvsn function implemented in the vsn package v3.72.0 (68) before any statistical testing. Gene expression analyses were performed with coding probes mapped to ENSEMBL gene IDs. QC was performed by graphical tools with the Uniform Manifold Approximation and Projection (UMAP) algorithm (M3C package v1.26.0 (69)). Differential expression (DE) analyses were performed with the limma algorithm (70) implemented in the piano package v2.20.0 (71). DE analyses were performed with pseudo-time contrasts (ND vs. sAAb+, sAAb+ vs. mAAb+, mAAb+ vs. T1D) and relative to control (sAAb+ vs. ND, mAAb+ vs. ND, T1D vs. ND) for INS+CD3-islets. A similar pseudo-time comparison was performed with INS+CD3+ islets for mAAb+ vs. T1D. In addition, INS+CD3+ islets from mAAb+ and T1D groups were compared to control (INS+CD3-ND) and to each corresponding non-infiltrated islet group. DE genes (DEGs) were defined as genes with an absolute logFC>0.5 and an adjusted *p*<0.001 unless otherwise stated. Over-representation (ORA) and gene set enrichment analyses (GSEA) with each DEG set were performed with the clusterProfiler package v4.12.6 (72–75) and included biological processes (BP) gene ontology (GO) as well as KEGG pathway enrichments. GSEA was run with an adjusted *p*<0.01. Supplementary tables are provided for all enrichment analyses performed across each contrast, as summarized in Table S2. DE analyses and GSEA were visualized with the graphical tools of clusterProfiler, ComplexHeatmap package v2.20.0 (76,77), aPEAR v1.0.0 (78), ggradar v0.2 (https://github.com/ricardo-bion/ggradar), and custom R scripts utilizing functions within those packages. The complete analysis pipeline and R code are posted to the GitHub repository at https://github.com/AlexandraCuaycal/laser_capture_microdissected_islet_transcriptomics. We also created an online tool to visualize normalized islet gene expression across the four clinical groups (https://alexandracuaycal.shinyapps.io/shiny_apps/).

### miRNA-target interactions database mining

DE miRNA genes in sAAb+ and mAAb+ islets (compared to ND) were interrogated in two different databases (prediction-based miRDB (24) and experimentally validated miRTarBase (25)) for their target interactions. miRNA targets were downloaded from both databases during the period of May 5-13, 2025, and data processed in RStudio v2024.04.0. We retrieved miRNA targets from both the 5p and 3p miRNA strands and analyzed them together. We performed miRNA target enrichment analysis in the DEGs from both sAAb+ and mAAb+ islets (vs. ND) with a Fisher’s exact test and corrected for multiple testing with the Benjamini-Hochberg (BH). We considered a significant enrichment of miRNA targets among the DEGs in sAAb+ and mAAb+ islets as those with adjusted p-value≤0.001 and OR>1. When relevant, we interrogated miRNAs in the Human microRNA Disease Database (HMDD). As a control, we also examined the miRNA target enrichments from miRNAs that were not significantly DE across all three clinical groups (sAAb, mAAb, T1D vs. ND) and with expected number of targets under 100 in both databases. We considered a non-significant enrichment after a Fisher’s exact test as those miRNAs with adj. p-values>0.05 for at least one database.

### Statistical analysis of gene and protein expression across groups

Gene expression plots across clinical and/or histological phenotypes were generated with variance-stabilized expression intensities as described above, and statistical testing was performed with a one-way analysis of variance (ANOVA) followed by Tukey’s post-hoc test. Statistical testing of corrected (c)MFI values across clinical and histological phenotypes was performed with the non-parametric Wilcoxon test. Correlation analyses were performed with Pearson’s. Multiple testing correction was performed with the BH method. Statistical significance was defined by adjusted *p*≤0.05.

### Study approval

nPOD operates under approval from the UF IRB. Organ and tissue recoveries are performed after informed consent has been obtained from the donors’ legal representatives (79). This study was performed under UF IRB201400796.

## Data availability

The microarray dataset can be accessed through the NCBI Gene Expression Omnibus (GEO) with accession number: GSE284772. The SST-RMA normalized microarray signal expression matrix can be downloaded from https://doi.org/10.5281/zenodo.14537115. The R code and analysis pipeline are posted to the GitHub repository at: https://github.com/AlexandraCuaycal/laser_capture_microdissected_islet_transcriptomics.

The supplementary data tables are available in the Supplementary Tables File.

## Author contributions

AEC performed the mIF and transcriptomics data curation, normalization, bioinformatics, statistical analyses, and wrote the original draft. EAB conducted islet microdissection, RNA isolation and analysis, mIF staining, and confocal imaging. SS and JC performed confocal imaging and mIF image analysis. NIL conducted microarray analysis. LAB performed miRNA-target data mining and curation. EAP provided guidance on data interpretation. SG wrote custom scripts for IF data retrieval. MAA provided expertise on experimental design and data interpretation. MCT, ICG, and CEM acquired funding, conceived of the study, supervised all aspects, and edited the manuscript. All authors have read and approved submission of the manuscript.

## Acknowledgements

The authors thank the nPOD donors and their families for organ donation to research. We also thank the nPOD Organ Processing and Pathology Core Staff at the University of Florida for providing OCT sections. The studies were accomplished with support from the UF Center for Immunology and Transplantation. The content and views expressed are the responsibility of the authors and do not necessarily reflect the official view of nPOD. Organ Procurement Organizations (OPO) partnering with nPOD to provide research resources are listed at https://npod.org/for-partners/npod-partners.

