## Supplemental Figures for "Mitochondrial and protein homeostasis pathways are transcriptionally impaired in islets during type 1 diabetes pathogenesis"

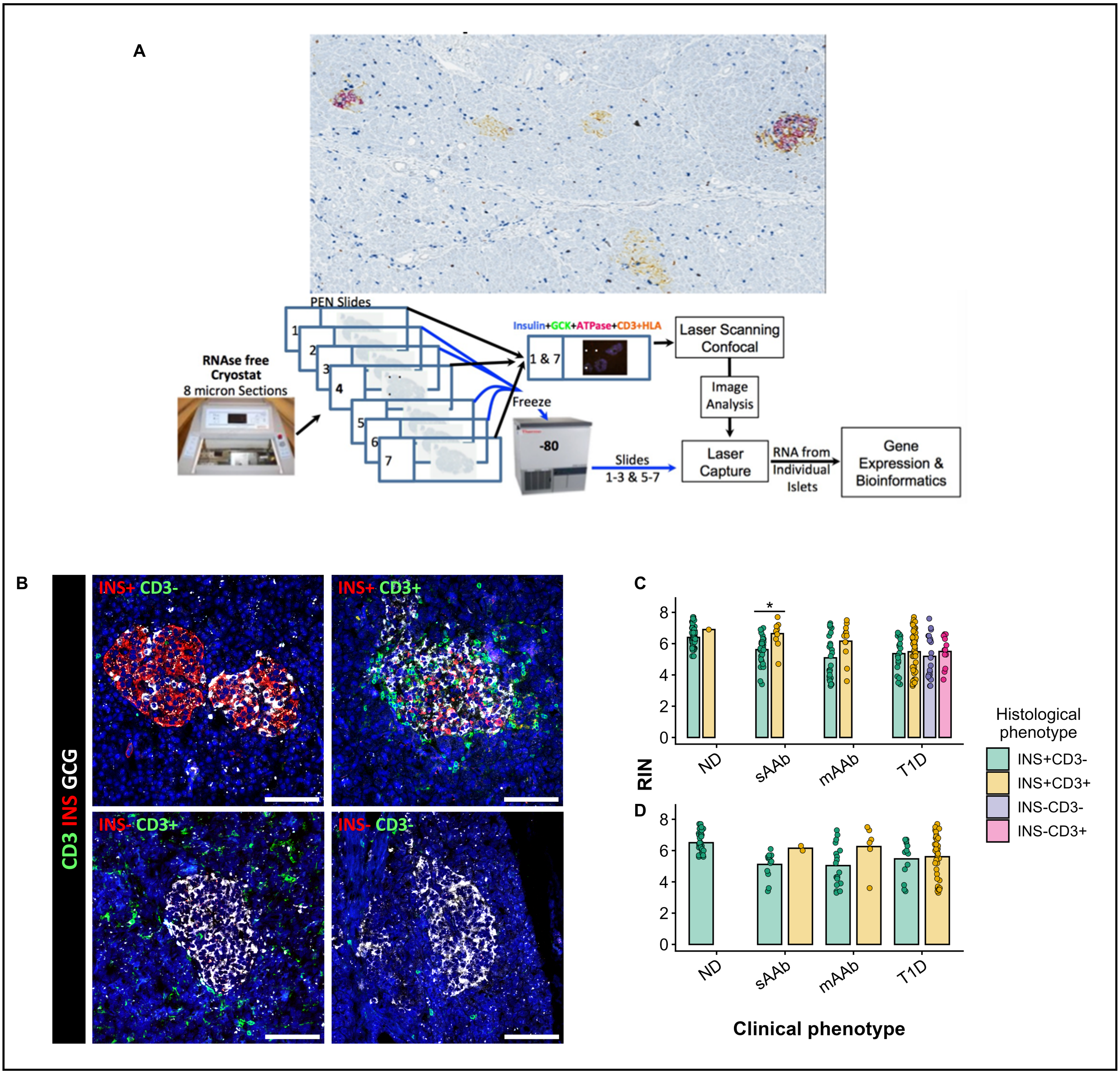


**Supplemental Figure 1. Experimental design. (A)** Schematic representation of pancreas tissue section acquisition and allocation for laser microdissection (LMD) and multiplex immunofluorescence (mIF) and subsequent analysis. **(B)** Histological phenotypes of islets assigned based on the insulin (INS) and CD3 stains. Size bar: 100 µm. **(C)** RNA integrity numbers (RIN) for the 260 islets across all clinical and histological phenotypes and for the final islet set post-QC **(D)**. Adjusted *p*: $*p<0.05$, $**p<0.01$, $***p<0.001$, $****p<0.0001$.

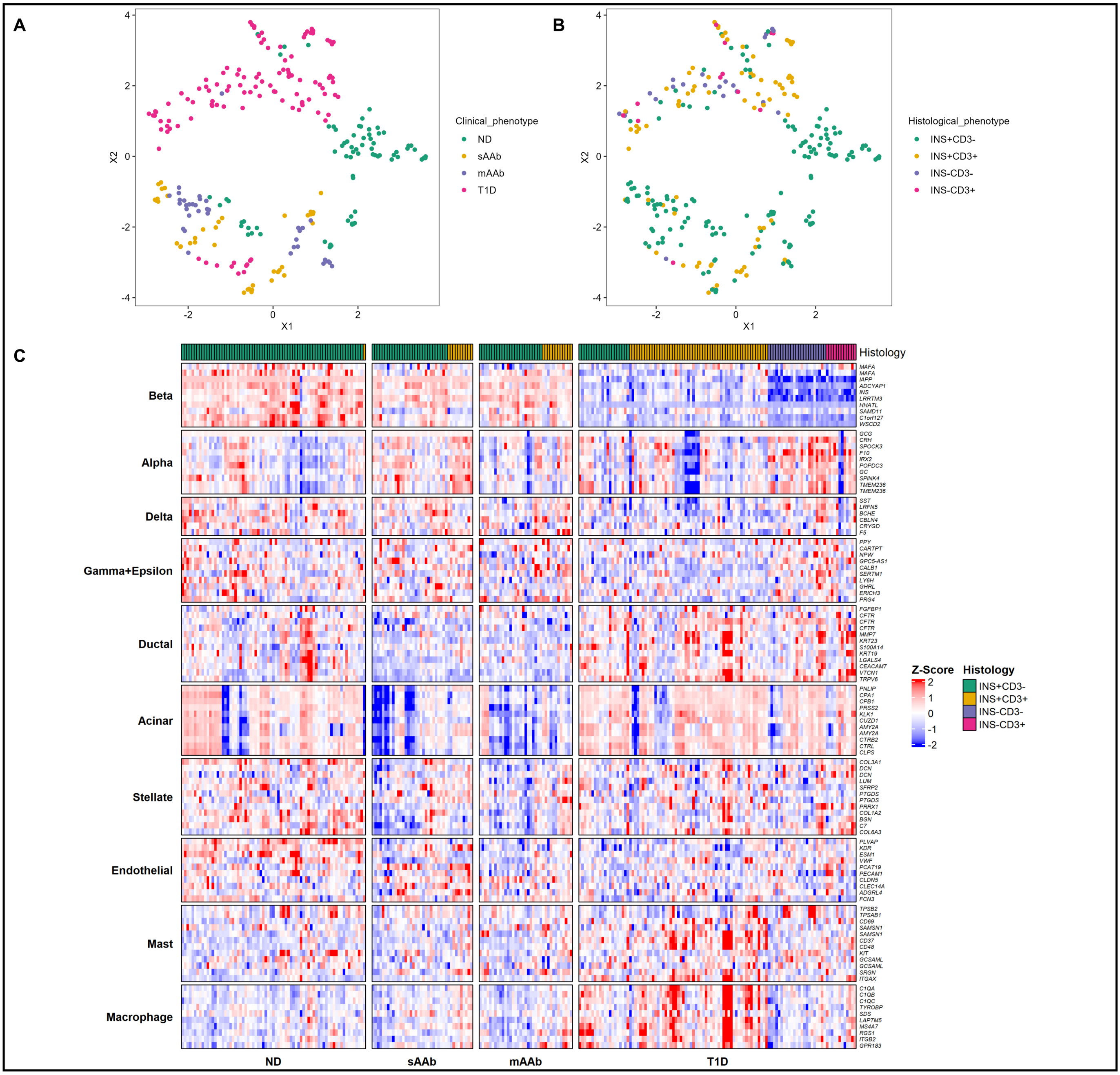


**Supplemental Figure 2. Exploratory analysis of 260 islets.** UMAP representations of islets colored by **(A)** clinical phenotype or **(B)** histological phenotype. **(C)** Heatmap of islet cell-type specific markers. Rows represent individual genes. Cluster of rows represent individual cell types. Columns represent individual islets. Cluster of columns represent clinical phenotypes. The heatmap top annotation is colored by histological phenotype of islets.

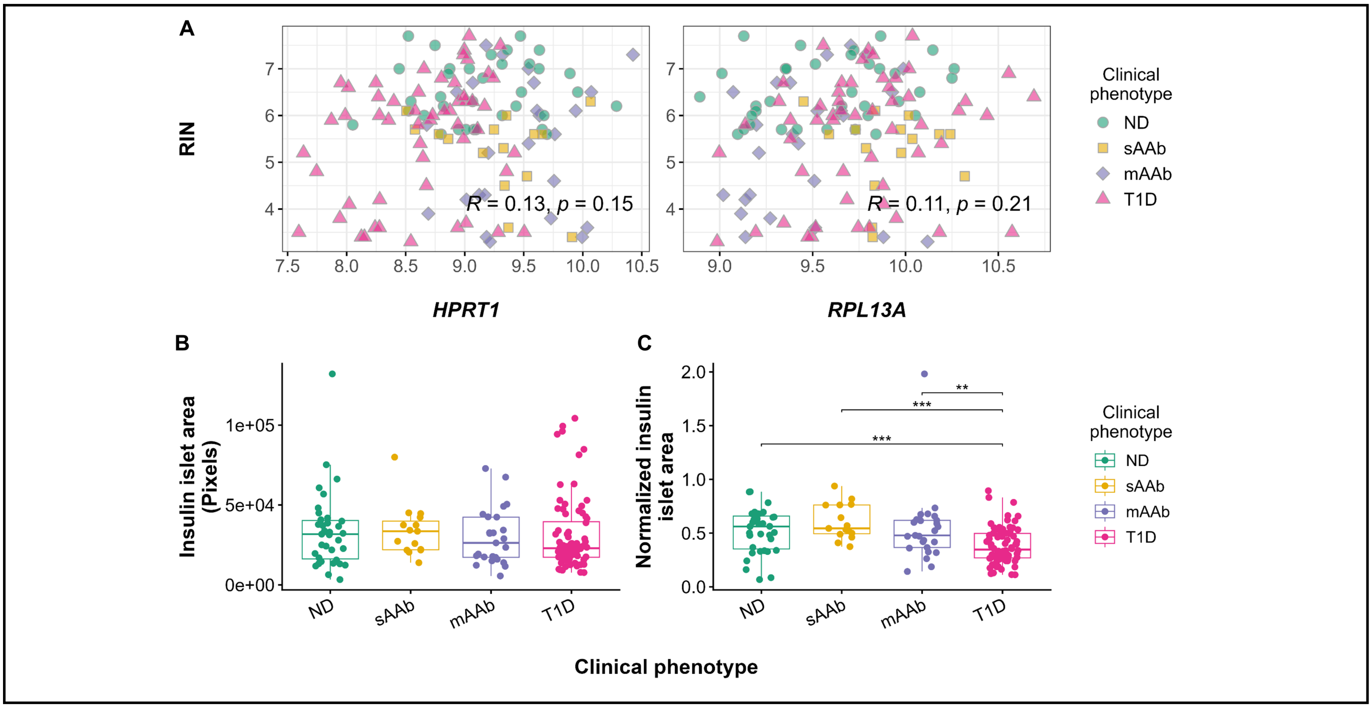


**Supplemental Figure 3. Metadata parameters of final islet set.** Pearson correlation analyses with housekeeping gene expression and RNA integrity numbers (RIN) **(A)**. Islet insulin area from insulin cMeanFI is shown as pixels **(B)** and normalized to islet cut area **(C)**. Statistical significance of Pearson correlations was considered with a $p<0.05$. Adjusted p: ∗p<0.05, ∗∗p<0.01, ∗∗∗p<0.001, ∗∗∗∗p<0.0001.

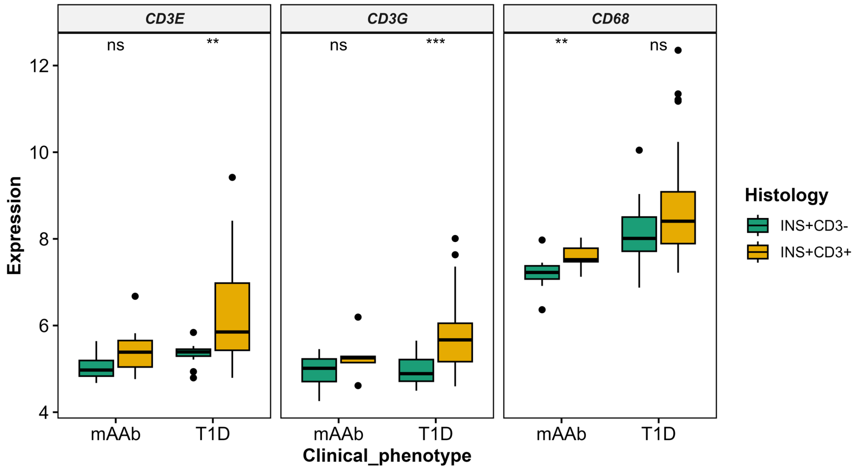


**Supplemental Figure 4. Gene expression of immune cell markers is increased after T-cell infiltration.** Boxplots of gene expression of *CD3E, CD3G* (T-cell markers), and *CD68* (macrophage marker), respectively. Gene expression is shown for both INS+CD3- and INS+CD3+ mAAb and T1D islets. Pair-wise comparisons between histological phenotypes were performed. $p$: $*p<0.05$, $**p<0.01$, $***p<0.001$, $****p<0.0001$.

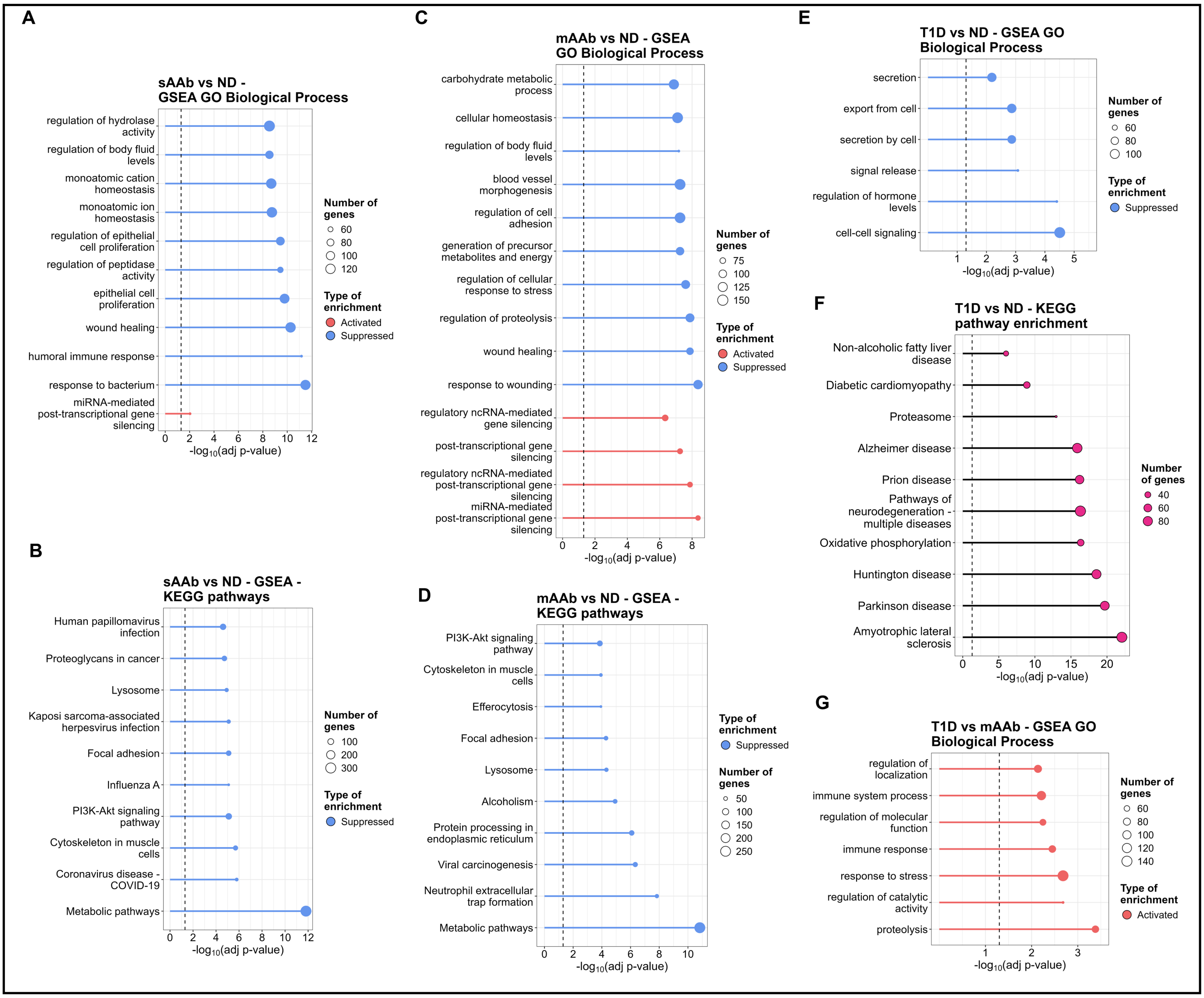


**Supplemental Figure 5. Enrichment analyses across INS+CD3- comparisons analyzed.** Gene set enrichment analysis (GSEA) plots show the top 10 activated and suppressed pathways from Gene Ontology Biological Process (GO-BP) and KEGG analyses for sAAb vs ND (**A** and **B**), mAAb vs ND (**C** and **D**), T1D vs ND (**E** and **F**), and T1D vs mAAb (**G**). **F** displays KEGG pathways over-representation analysis (ORA). Statistical significance for BPs and KEGG pathways in GSEA and ORA was considered with an adjusted $p<0.01$.

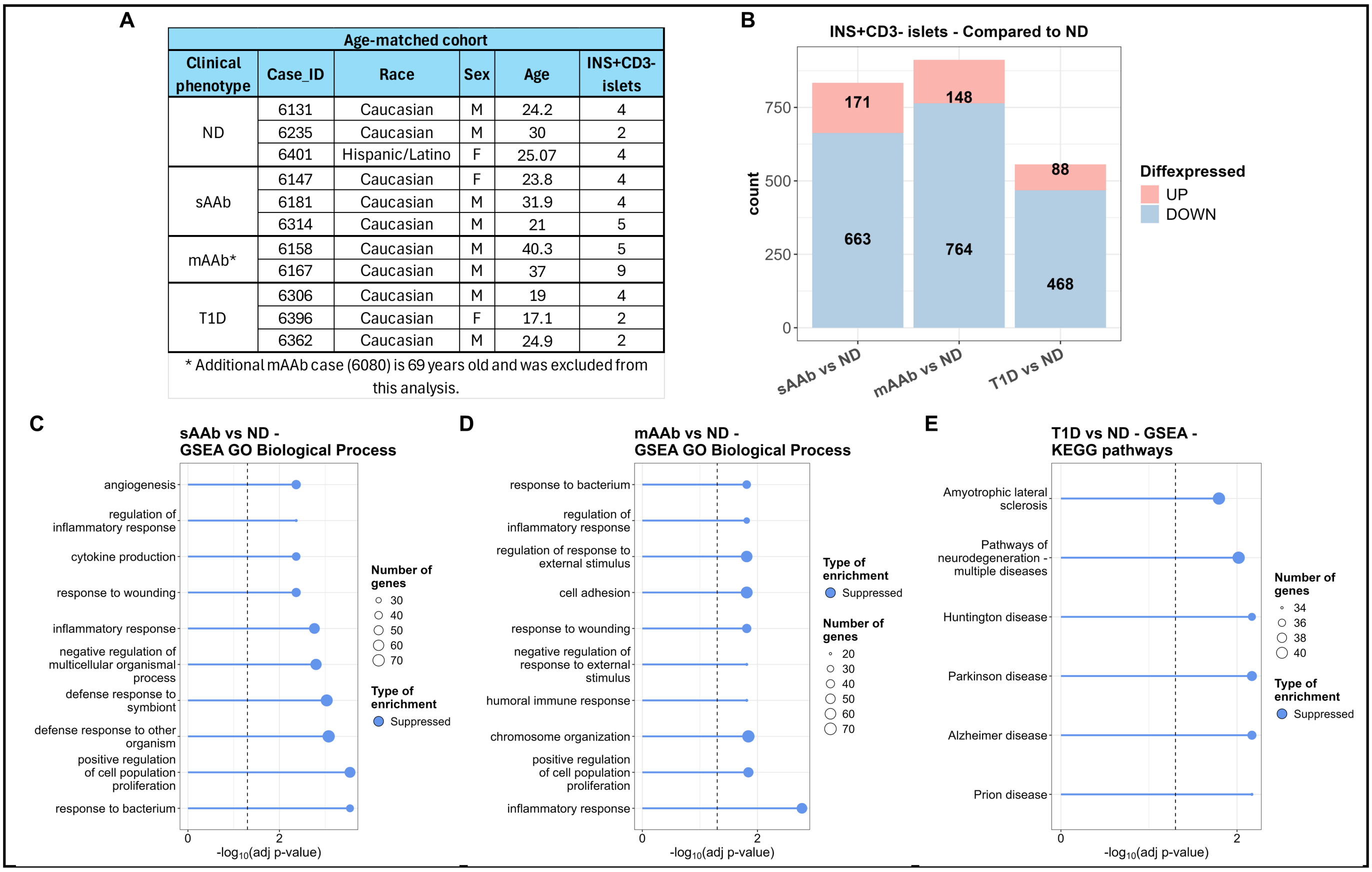

| Clinical phenotype | Case_ID | Race | Sex | Age | INS+CD3- islets |
|---|---|---|---|---|---|
| ND | 6131 | Caucasian | M | 24.2 | 4 |
| | 6235 | Caucasian | M | 30 | 2 |
| | 6401 | Hispanic/Latino | F | 25.07 | 4 |
| sAAb | 6147 | Caucasian | F | 23.8 | 4 |
| | 6181 | Caucasian | M | 31.9 | 4 |
| | 6314 | Caucasian | M | 21 | 5 |
| mAAb* | 6158 | Caucasian | M | 40.3 | 5 |
| | 6167 | Caucasian | M | 37 | 9 |
| T1D | 6306 | Caucasian | M | 19 | 4 |
| | 6396 | Caucasian | F | 17.1 | 2 |
| | 6362 | Caucasian | M | 24.9 | 2 |



**Supplemental Figure 6. Differential expression (DE) analyses of INS+CD3- islets in an age-matched donor subcohort.** Donors across clinical groups were age-matched, and corresponding islet counts are shown (**A**). Bar plots show the number of DEGs identified for each comparison analyzed (**B**). DEGs were defined as genes with an absolute logFC>0.5 and an adjusted *p*<0.001. Gene set enrichment analysis (GSEA) plots show the top 10 activated and suppressed gene ontology biological process (GO-BP) pathways for the sAAb vs ND (**C**) and mAAb vs ND (**D**) comparisons. GSEA of KEGG pathways is shown for the T1D vs ND comparison (**E**). Statistical significance for BPs and KEGG pathways in GSEA and ORA was considered with an adjusted *p*<0.01.

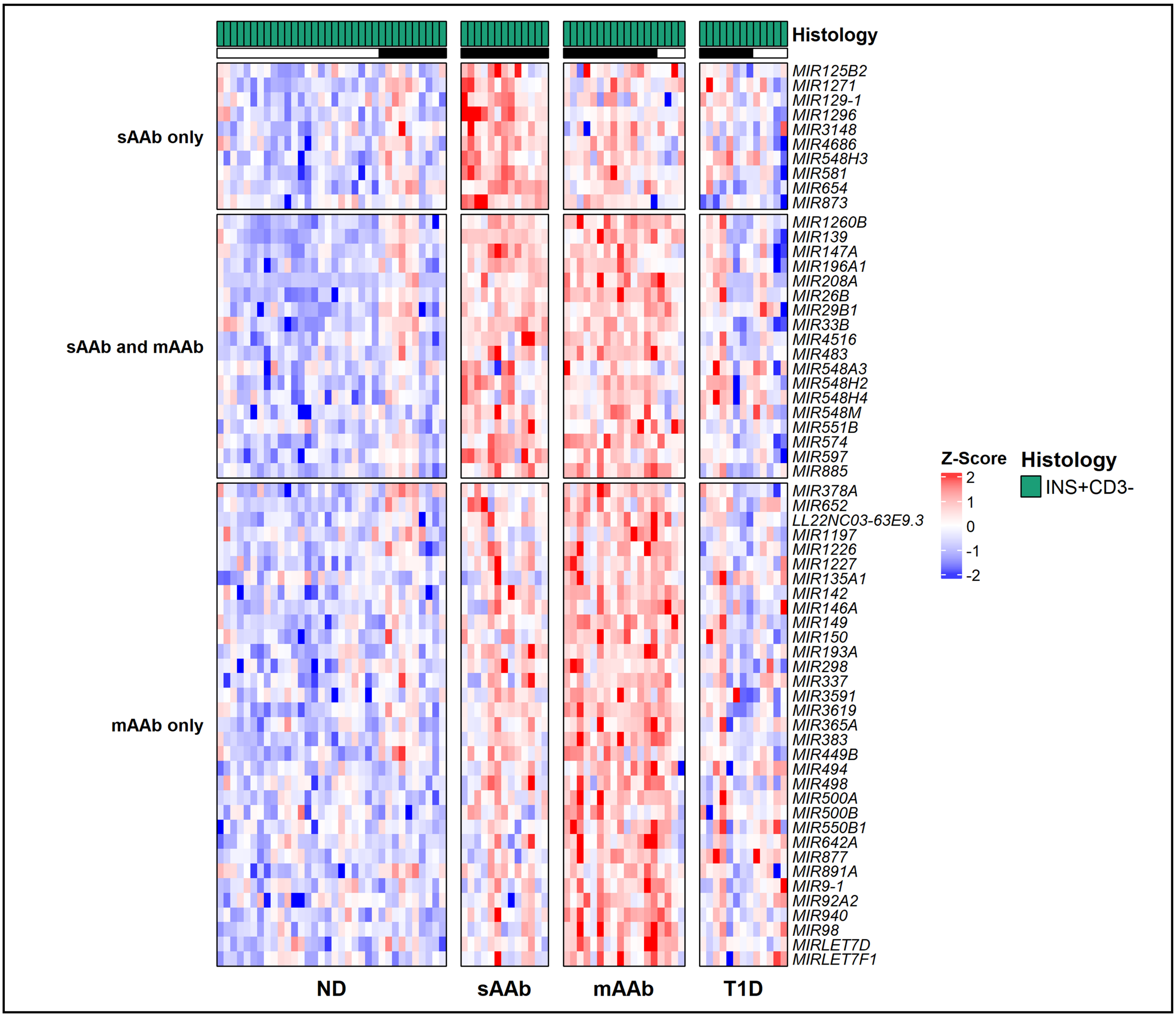


**Supplemental Figure 7. Heatmap of upregulated miRNA genes in INS+CD3- sAAb and mAAb vs ND islets.** Gene expression in the heatmap is presented as z-scores. The heatmap presents the gene expression across all clinical groups for miRNA genes identified as DE in sAAb and mAAb vs ND islets. Black line on top indicates islets from age-matched subcohort.

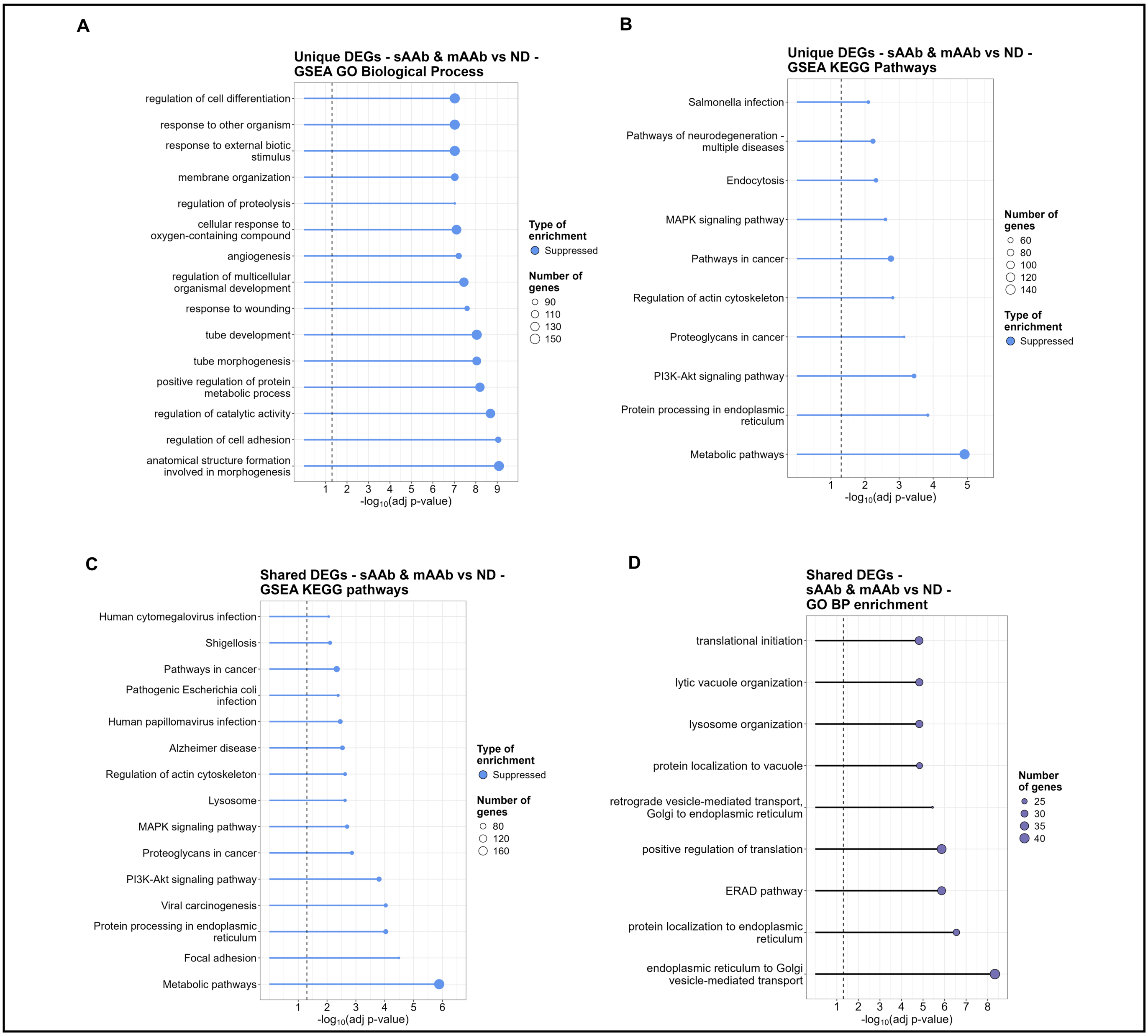


**Supplemental Figure 8. Enrichment analyses for common DEGs in INS+CD3- islets from sAAb and mAAb vs ND comparisons.** Gene set enrichment analyses (GSEA) plots show the top 15 activated and suppressed pathways from Gene Ontology Biological Process (GO-BP) and KEGG analyses. Pathways are shown for DEGs common to the sAAb vs ND and mAAb vs ND comparisons but not shared with T1D vs ND (**A** and **B**), and for DEGs common to sAAb vs ND, mAAb vs ND, and T1D vs ND comparisons (**C** and **D**). **D** shows over-representation analysis (ORA) for GO-BP. Statistical significance for BPs and KEGG pathways in GSEA and ORA was considered with an adjusted $p<0.01$.

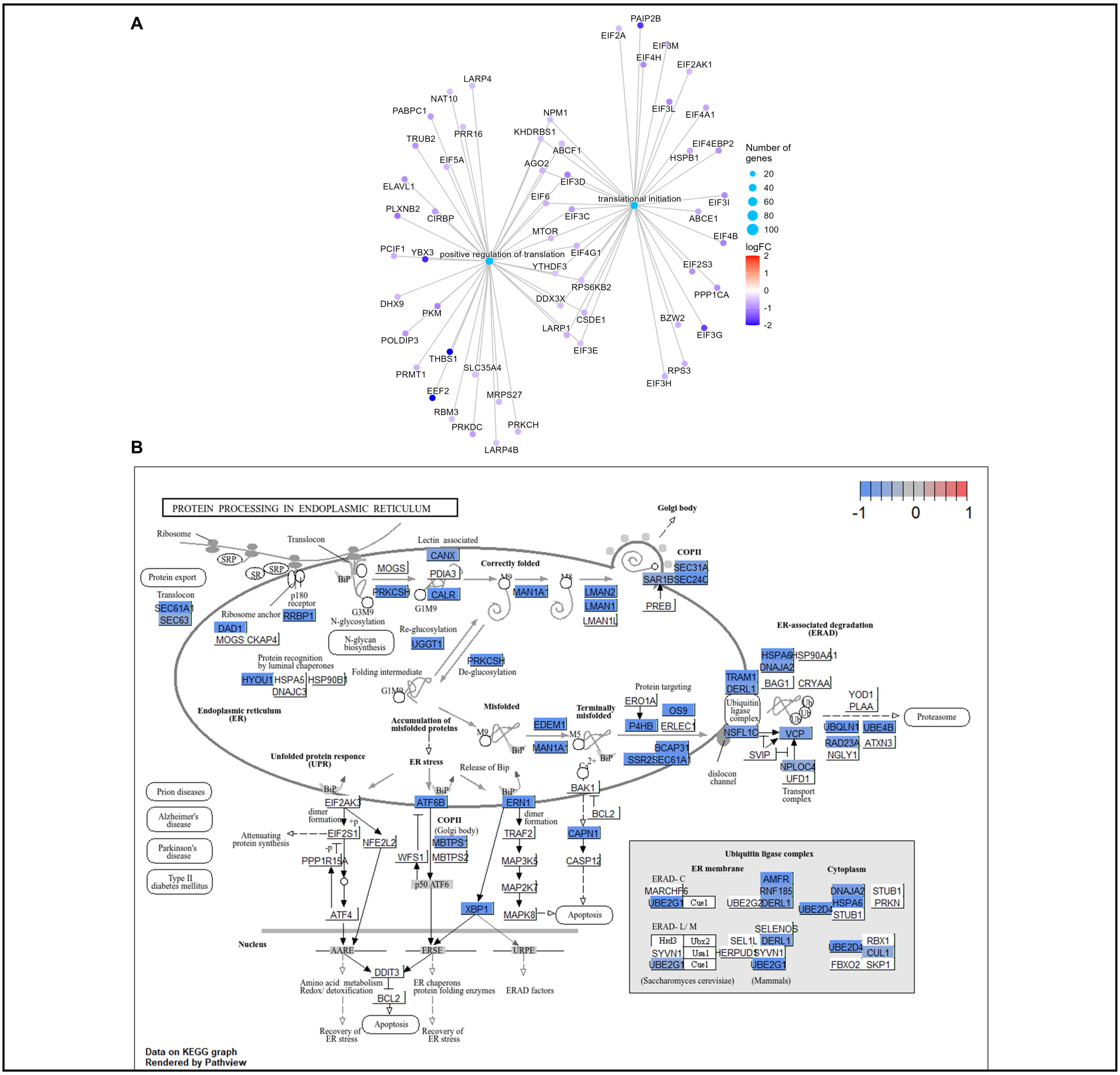

**Supplemental Figure 9. Downregulation of translation and ER protein-processing genes in sAAb and mAAb INS⁺CD3⁻ islets compared to ND.** Gene-concept network plot showing differential gene expression of translation pathways **(A)** and the KEGG pathway map for "Protein processing in the endoplasmic reticulum" **(B)**. Gene expression in A and B is shown as log2FC values for the mAAb vs ND comparison.

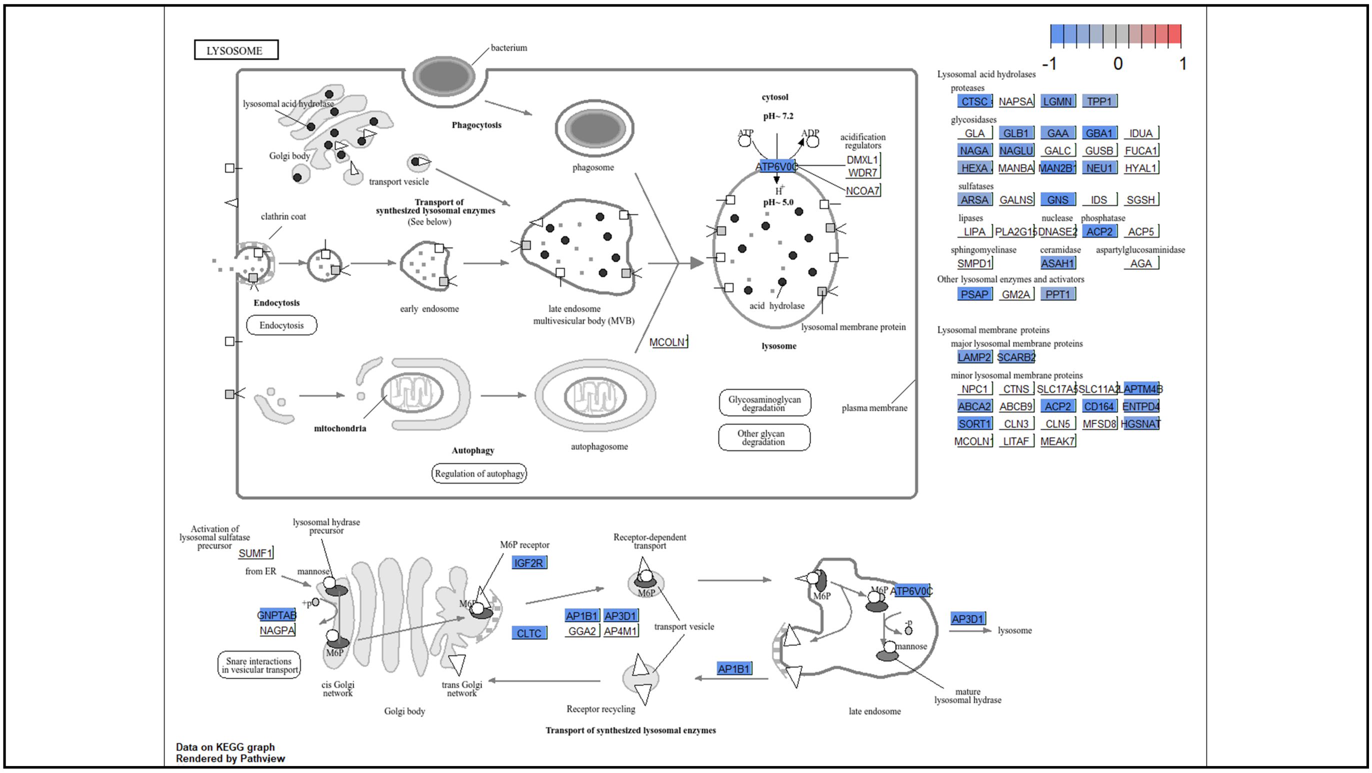


**Supplemental Figure 10. Lysosome KEGG pathway map for the DEGs in INS+CD3- islets from the sAAb and mAAb vs ND comparison.** Gene expression in the pathway map is shown as log2FC values for the mAAb vs ND comparison.

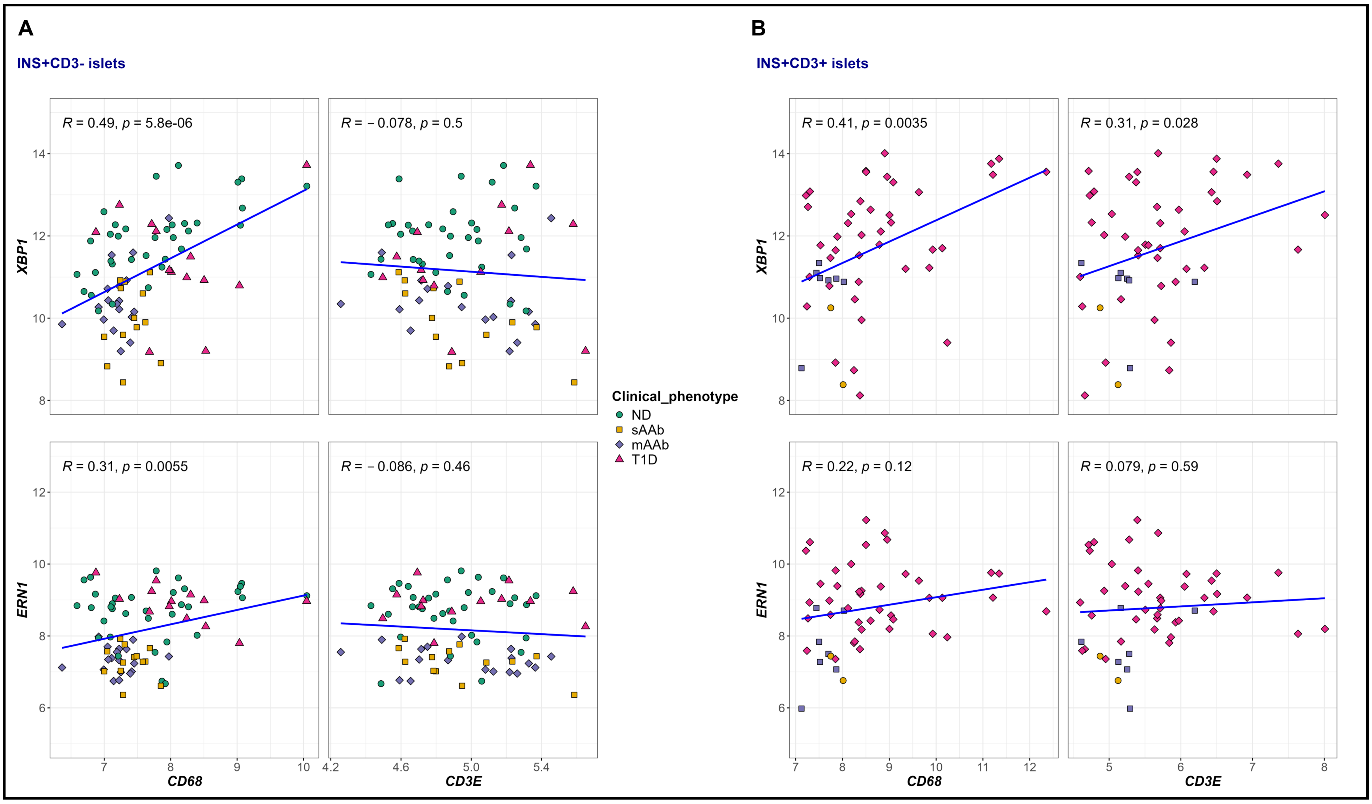


**Supplemental Figure 11. Pearson correlation plots for ER stress vs immune cell markers across clinical groups.** Correlation plots are shown for both INS+CD3- **(A)** and INS+CD3+ **(B)** islets. Statistical significance of Pearson correlations was considered with a *p*<0.05.

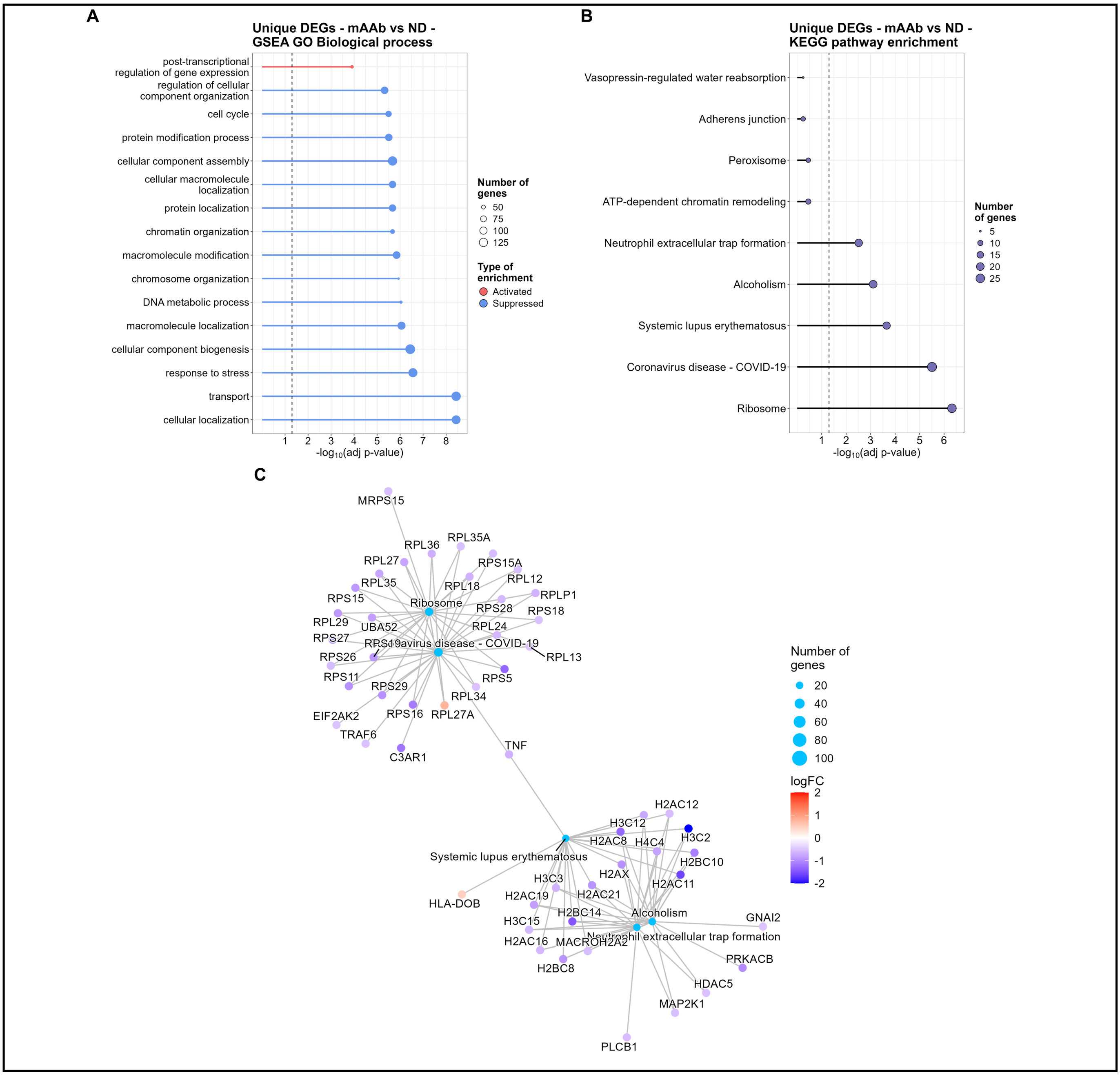


**Supplemental Figure 12. Enrichment analyses for unique DEGs in INS+CD3- islets from mAAb vs ND comparison.** Gene set enrichment analysis (GSEA) plot show the top 15 activated and suppressed pathways from Gene Ontology Biological Process (GO-BP) **(A).** Over-representation analysis (ORA) plot shows the top 15 KEGG pathways (**B**). Enrichment analyses were done for DEGs unique to the mAAb vs ND comparison. The gene-concept network plot for ribosomal-related pathways is shown in **C**. Statistical significance for BPs and KEGG pathways in GSEA and ORA was considered with an adjusted *p*<0.01.

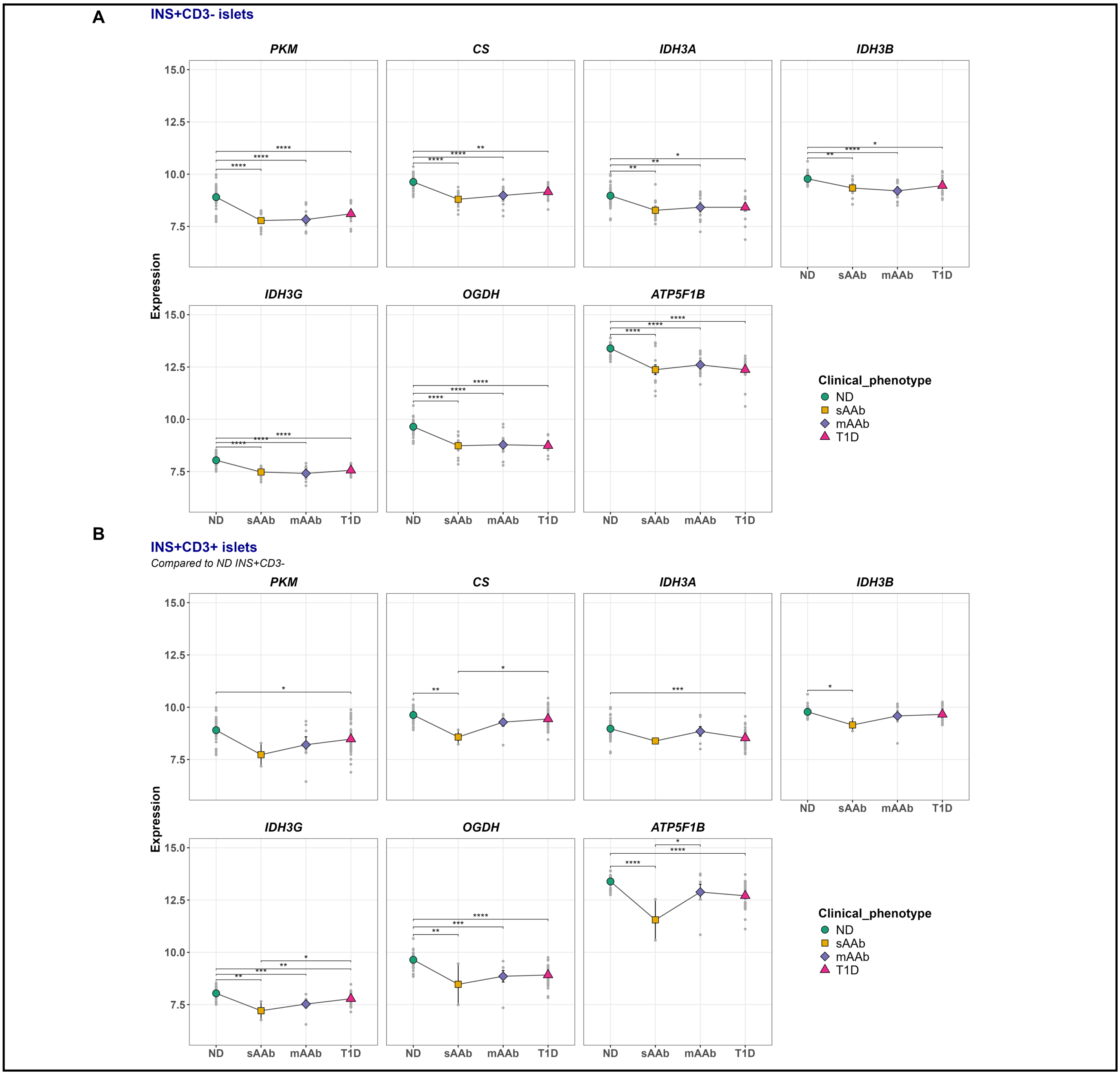


**Supplemental Figure 13. Gene expression of glycolytic and TCA cycle rate-liming enzymes that are differentially expressed in INS+CD3- islets from the mAAb vs ND comparison.** Gene expression plots show average islet expression color-coded by clinical group, with single-islet expression shown in gray, for non-infiltrated INS+CD3- islets (**A**) and infiltrated INS+CD3+ islets (**B**). The ND group in B corresponds to INS+CD3- islets. Adjusted *p*: $*p<0.05$, $**p<0.01$, $***p<0.001$, $****p<0.0001$.

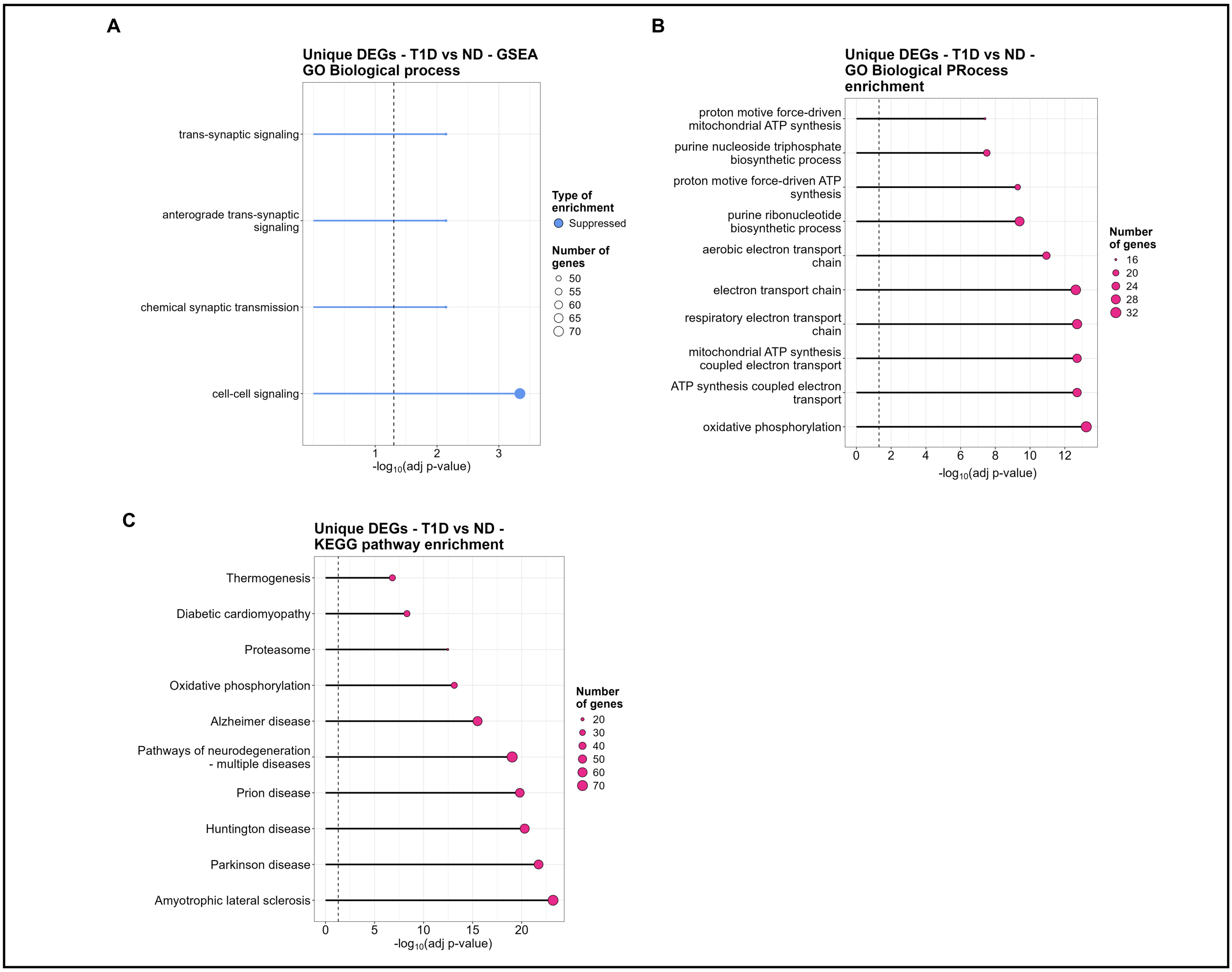


**Supplemental Figure 14. Enrichment analyses for unique DEGs in INS+CD3- islets from T1D vs ND comparison.** Gene set enrichment analysis (GSEA) plot show the top 10 activated and suppressed pathways from Gene Ontology Biological Process (GO-BP) **(A).** Over-representation analysis (ORA) for GO-BP (**B**) and KEGG pathways (**C**). All enrichment analyses were run with DEGs unique to the T1D vs ND comparison (INS+CD3-). Statistical significance for BPs and KEGG pathways in GSEA and ORA was considered with an adjusted *p*<0.01.

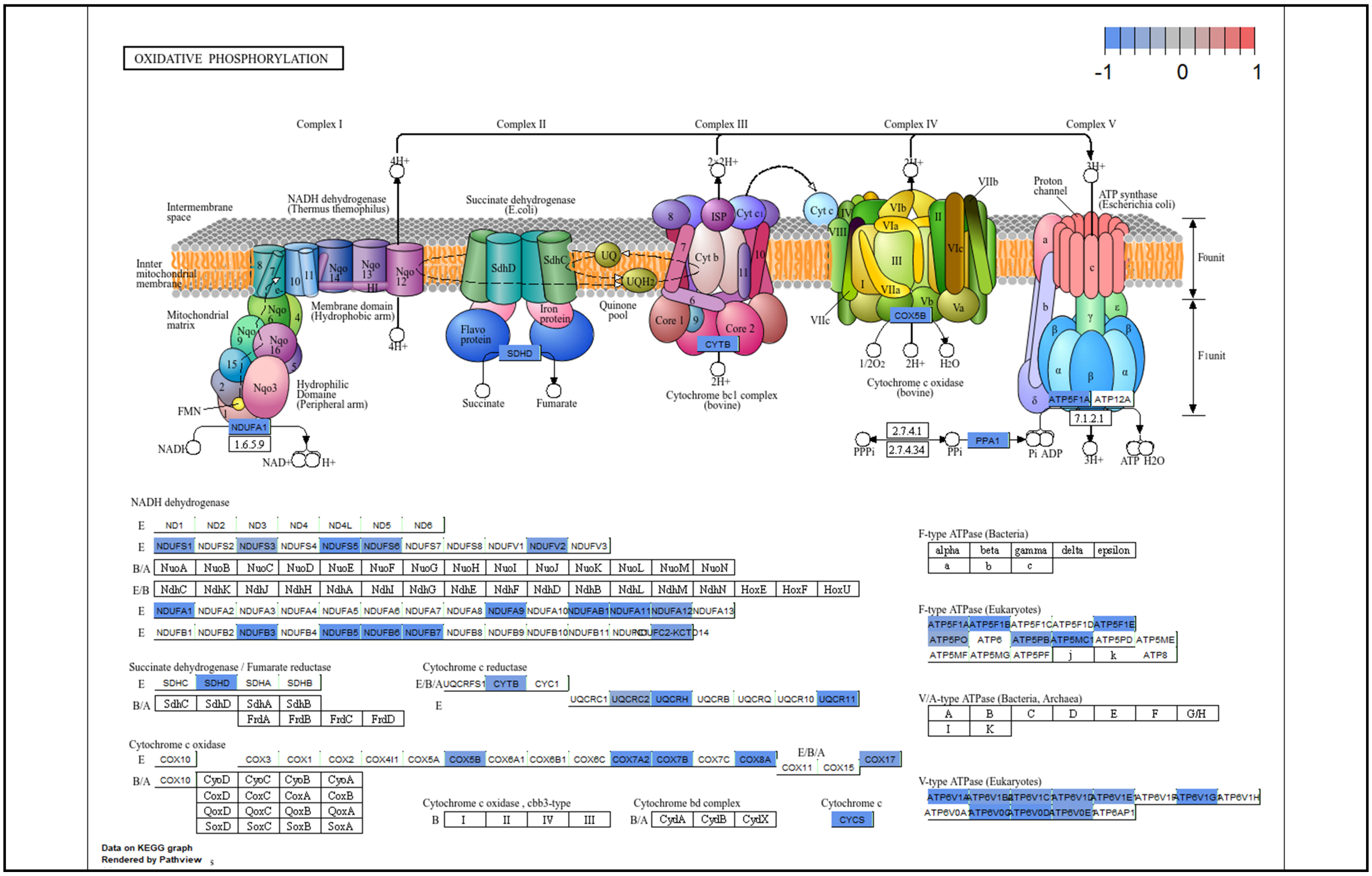


**Supplemental Figure 15. Oxidative phosphorylation is downregulated in INS+CD3- islets from T1D vs ND comparison.** The figure shows the KEGG pathway map with gene expression colored by logFC values from the T1D vs ND comparison.

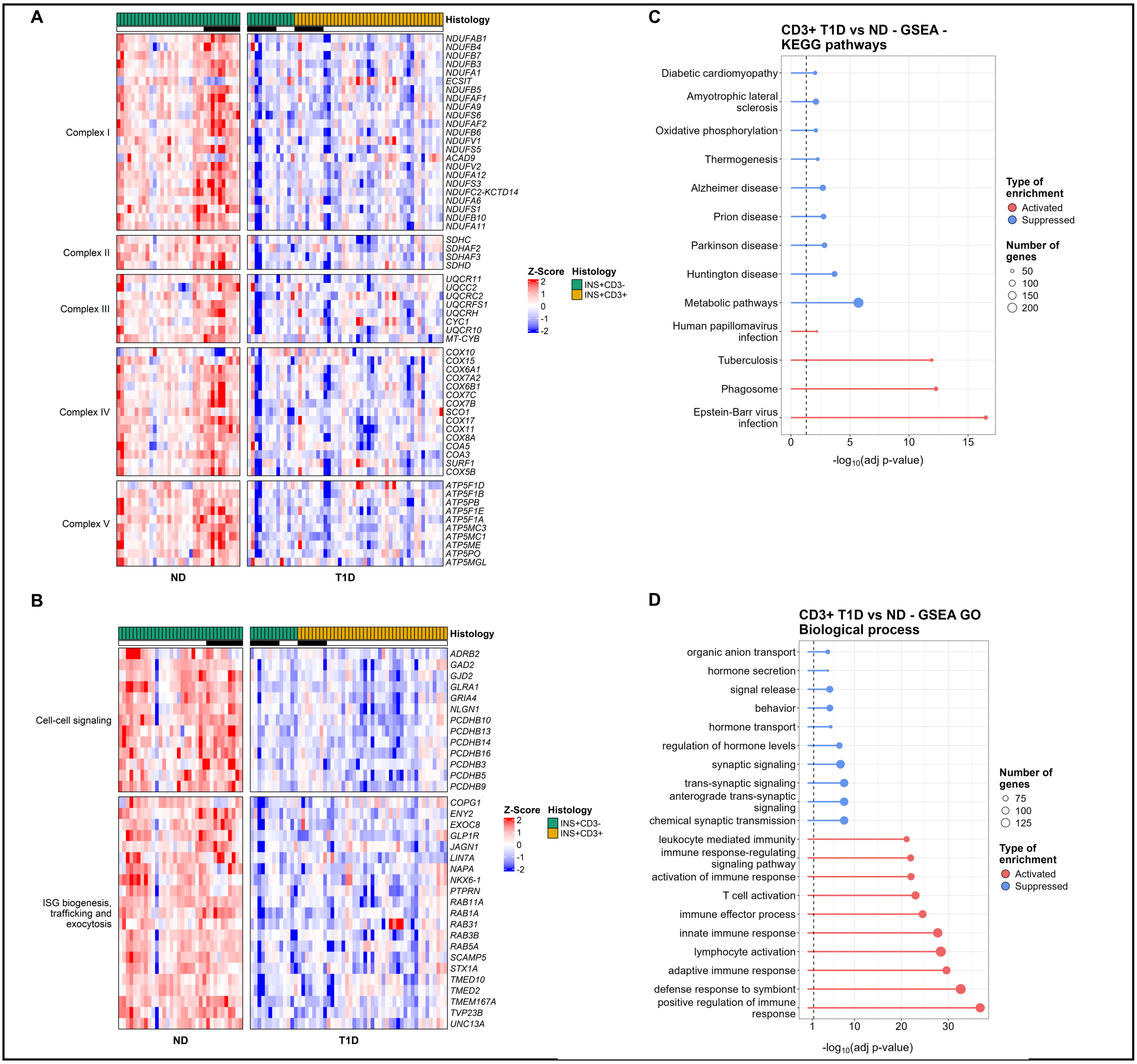


**Supplemental Figure 16. Dysregulated pathways in INS+CD3- T1D islets remained similarly altered in INS+CD3+ T1D islets compared to ND.** Heatmaps show DEGs across electron transport chain (ETC) complexes (**A**) and insulin release-related pathways (**B**). Black line on top of heatmaps indicates islets from age-matched subcohort. Gene set enrichment analyses (GSEA) show the top 10 activated and suppressed KEGG (**C**) and gene ontology biological process (GO-BP) pathways (**D**). GSEA in C and D were run with DEGs in infiltrated (INS+CD3+) T1D vs ND islets. Statistical significance for BPs and KEGG pathways in GSEA and ORA was considered with an adjusted $p<0.01$. Gene expression in the heatmap is presented as z-scores.

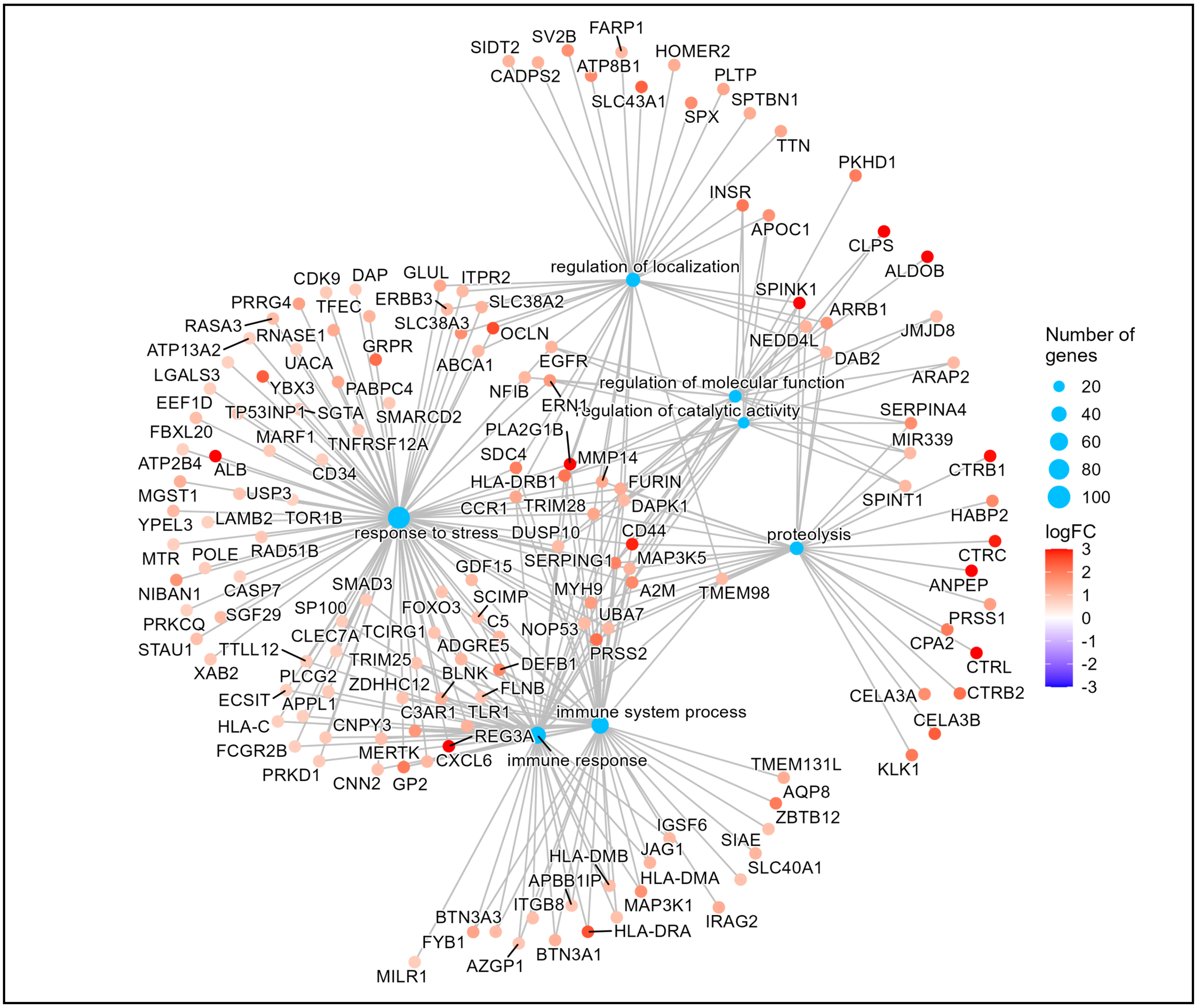


**Supplemental Figure 17. Activated pathways in the pseudo-transition from mAAb to T1D in INS+CD3- islets.** Gene-concept plot showing all significantly enriched pathways in the T1D vs mAAb comparison. Gene expression is shown as logFC values from the T1D vs mAAb comparison.
